# A powdery mildew core effector protein targets host endosome tethering complexes HOPS and CORVET

**DOI:** 10.1101/2024.07.17.603885

**Authors:** Björn Sabelleck, Sohini Deb, Pietro D. Spanu, Hans Thordal-Christensen, Ralph Panstruga

## Abstract

Powdery mildew fungi are serious pathogens of many plant species. Their genomes encode extensive repertoires of secreted effector proteins that suppress host immunity. Here, we revised and analyzed the candidate secreted effector protein (CSEP) effectome of the barley powdery mildew fungus, *Blumeria hordei* (*Bh*) and identified seven CSEPs that are broadly conserved in powdery mildew fungal species, rendering them core effectors of these phytopathogens. We show that one of these, CSEP0214, interacts with the barley VPS18 protein, a core component of the CORVET and HOPS endosomal tethering complexes, mediating fusions of early endosomes and multivesicular bodies with the central vacuole, respectively. Overexpression of CSEP0214 and knockdown of either *VPS18*, HOPS-specific *VPS41* or CORVET-specific *VPS8* caused a block of the vacuolar pathway and the accumulation of the fluorescent vacuolar marker protein (SP)-RFP-AFVY in the endoplasmic reticulum. Co-immunoprecipitation and yeast two-hybrid experiments suggest that CSEP0214 blocks the interaction of VPS18 and VPS16, which are both core components of CORVET as well as HOPS. Additionally, expression of CSEP0214 blocked the hypersensitive cell death response associated with resistance gene-mediated immunity in barley, indicating that endomembrane traffic is required for this process. It also prevented callose deposition in cell wall appositions at attack sites and encasements of fungal infection structures. Our results indicate that this powdery mildew core effector is an essential immunity suppressor.

**One sentence summary:** The *Blumeria hordei* effector protein CSEP0214 interacts with barley VPS18, a core component of the CORVET and HOPS endosomal tethering complexes, thereby interfering with host endomembrane trafficking.

## Introduction

The powdery mildew fungi are obligate biotrophic pathogens belonging to the phylum Ascomycota and comprise ∼900 species, which can infect nearly 10,000 species of angiosperm plants (Braun and Cook, 2012). In the course of the infection powdery mildew fungi secrete up to hundreds of effector proteins into the apoplastic space and the host cytosol to suppress plant defence responses and to manipulate the host cell metabolism for the benefit of the pathogen (Barsoum et al., 2019).

The powdery mildew pathogen, *Blumeria hordei* (*Bh*, formerly known as *B. graminis* f.sp. *hordei* (Liu et al., 2021)), causing disease on barley (*Hordeum vulgare*), can lead to significant yield losses in the field. Its unusually large genome codes for around 800 secreted proteins many of which are thought to act as effectors to promote pathogen virulence (Frantzeskakis et al., 2018). Of these, only few have been functionally characterized so far (Aguilar et al., 2016; Ahmed et al., 2015; Ahmed et al., 2016; Li et al., 2021; Liao et al., 2023; Pennington et al., 2019; Pliego et al., 2013; Schmidt et al., 2014; Yuan et al., 2021; Zhang et al., 2012).

Upon recognition of fungal molecular structures, the plant immune system is activated in discrete steps (Jones and Dangl, 2006), involving pattern-triggered immunity (PTI) and effector-triggered immunity (ETI) (Thordal-Christensen, 2020). PTI comprises complex transcriptional reprogramming and cellular responses, including the deposition of cell wall appositions (papillae) at the sites of attack and the formation of encasements in the form of cell wall extensions that enclose pathogen structures (haustoria) invading the plant cell (Underwood, 2012). As a countermeasure, the pathogen secretes effector proteins into the plant cytosol and apoplastic space to dampen and/or delay PTI.

Several *Arabidopsis thaliana* proteins are important for preinvasive immunity towards the non-adapted pathogen *Bh*, including the PENETRATION (PEN) proteins (Underwood and Somerville, 2008). The target membrane-associated soluble *N*-ethylmaleimide-sensitive-factor attachment receptor (t-SNARE, also referred to as syntaxin) SYP121 (synonym PEN1) is a component of an evolutionarily conserved defence pathway that relies on secretory processes (Rubiato et al., 2022). PEN1 and its orthologue in barley, ROR2, are required for papilla and encasement formation, probably by conferring fusion of multivesicular bodies (MVBs) to the plasma membrane (PM) at the site of fungal attack. Hereby MVB intraluminal vesicles are secreted as extracellular vesicles (EVs) towards papillae and encasements, which are labeled with PEN1 or ROR2 (Assaad et al., 2004; Böhlenius et al., 2010; Meyer et al., 2009; Nielsen et al., 2012; Nielsen et al., 2017; Rubiato et al., 2022; Ruf et al., 2022; Collins et al., 2003).

Transport along the endomembrane trafficking pathway requires membrane fusion events involving early and late endosomes. In addition to SNARE proteins, this process needs endosomal tethering factors, e.g. the Class C ‘core vacuole/endosome tethering’ (CORVET) complex that interacts with Rab5 GTPases, as well as the ‘homotypic fusion and protein-sorting’ (HOPS) complex interacting with Rab7 GTPases (Takemoto et al., 2018). The CORVET and HOPS complexes share the Class C vacuolar protein sorting (Vps) complex, made up of VPS11, VPS16, VPS18, and VPS33. In addition, CORVET involves VPS3 and VPS8, while HOPS involves VPS39 and VPS41 (Peterson and Emr, 2001; Rieder and Emr, 1997). Maturation of early endosomes to late endosomes, which eventually fuse with the vacuole, requires switches from the CORVET to the HOPS complex and from Rab5 to Rab7 (Rink et al., 2005).

Here we show that the *Bh* candidate effector CSEP0214, a highly conserved effector protein within the powdery mildew fungal lineage, interacts with the CORVET and HOPS complex protein, VPS18. Interaction occurs via, but not only, the conserved CFEM (common in fungal extracellular membrane) domain in CSEP0214 and the RING (really interesting new gene) domain of VPS18. We demonstrate that transient expression of *CSEP0214* in barley epidermal cells perturbs the endomembrane trafficking pathway to the vacuole. This results in the sequestration of marker proteins in the endoplasmic reticulum (ER), thereby preventing transport to their destinations. Importantly, the presence of CSEP0214 hampers encasement formation as well as the *Mla1*- and *Mla3*-mediated hypersensitive response (HR) in response to *Bh* challenge, confirming that endomembrane trafficking is essential for both PTI and ETI in the course of the barley-powdery mildew interaction. We finally observe that CSEP0214 prevents interaction between VPS18 and VPS16, another CORVET/HOPS protein, providing mechanistic insight into how CSEP0214 may suppress endomembrane trafficking.

## Results

### *In silico* analysis of the *B. hordei* secretome and the definition of a core effectome of powdery mildew fungi

The first in-depth analysis of the *Bh* ‘candidate secreted effector proteins’ (CSEPs) was performed more than a decade ago (Pedersen et al., 2012). Substantial advances during the last ten years in algorithms, prediction tools and computer power, the accessibility of new large data sets (e.g. transcriptomics and proteomics (Bindschedler et al., 2016)), and the release of an improved *Bh* genome assembly (Frantzeskakis et al., 2018), justify an updated survey of this group of proteins. Here, we aimed to provide a comprehensive summary and detailed *in silico* analysis of the entire *Bh* secretome, which includes an assignment of revised gene identifiers to CSEP numbers, the integration of published transcriptomics data, functional and structural predictions with InterPro, Phyre² and Alphafold2, and the use of general prediction tools like EffectorP and Localizer (see Materials and Methods for details).

The definition of effector proteins in eukaryotic pathogens (including fungi) is rather arbitrary and has been widely debated in the past, but typically assumes the presence of an amino-terminal signal peptide (SP) for secretion and the absence of detectable transmembrane domains (Sonah et al., 2016). Based on these criteria, we performed the present computational analysis with an updated list of 800 *Bh* proteins predicted to be secreted (Frantzeskakis et al., 2018) to reduce the risk of missing out on any potential effector candidates. According to the EffectorP 3.0 prediction, 527 of these 800 proteins were classified as *bona fide* effector candidates (328 cytoplasmic, 96 apoplastic and 103 cytoplasmic/apoplastic). This includes all known avirulence proteins of *Bh* and most CSEPs with identified plant targets. InterPro detected known domains in 266 of the 800 secreted proteins, whereby ribonuclease (IPR016191; 78 hits), protease/peptidase (various InterPro domains; 27 hits) and Egh16-like virulence factor (IPR021476; 23 hits) were the most prevalent assignments (Supplementary Table 1).

So far, only the core effectome of members of the *Blumeria* lineage, which infect grasses, was determined (Frantzeskakis et al., 2018); the core effectome of the entirety of monocot- and dicot-infecting powdery mildew fungi has not been studied. With this aim, we followed the bioinformatic strategy outlined in Figure 1. We used the predicted 800-protein secretome of *Bh*, which has one of the best annotated powdery mildew fungal genomes, as a query for sequence-based searches against 17 genomes of 15 additional powdery mildew fungi covering a broad range of the powdery mildew species diversity (Figure 1). Apart from the two grass-infecting *Blumeria* species, these fungi infect dicotyledonous host species, while no known powdery mildew fungi appear to infect non-grass monocots (Vaghefi et al., 2022). We performed TBLASTN searches with a relaxed e-value threshold of e^-10^ not to miss detection of any potential effector candidates in these *Bh* relatives. To account for incomplete and/or low quality genome sequences, we accepted candidate effectors that are present in the genomes of just 15 out of the 16 tested species. Based on these criteria, we detected 196 *Bh* sequences (Supplementary Table 2) that are conserved in the tested powdery mildew fungi. Among the remaining 604 proteins present in fewer than fifteen powdery mildew genomes, 30 were exclusively present in *Bh*, suggesting they are species-specific effector proteins of the barley pathogen. To eliminate secreted housekeeping proteins and other conserved secreted proteins from the list of the 196 secreted proteins conserved in at least 15 out of the 16 tested powdery mildew species, TBLASTN searches were carried out with the *Bh* versions of those as queries against a selection of genomes of non-phytopathogenic ascomycete fungi (*Aspergillus* spp., *Penicillium* spp., and *Saccharomyces* spp.). This resulted in 168 secreted proteins that can be considered as common fungal housekeeping proteins. A TBLASTN search was performed with the remaining 28 proteins against the proteomes of 23 other phytopathogenic Ascomycota (see legend of Figure 1) to explore a potential overlap of effector sets. This ultimately resulted in seven proteins that according to our analysis represent the conserved core effectome specific for the investigated powdery mildew fungi, while 21 proteins are also present in other plant-pathogenic fungi (Supplementary Table 2). One of the seven powdery mildew-specific core candidate effectors is CSEP0214 (*B. hordei* identifier BLGH_02334), a protein of 133 amino acids (including its amino-terminal SP), for which we provide a detailed *in silico* analysis, characterization of the interaction with its primary plant target, and an investigation of the cell biology and immunity-related phenotypes upon its expression in barley host cells.

**Figure 1.**
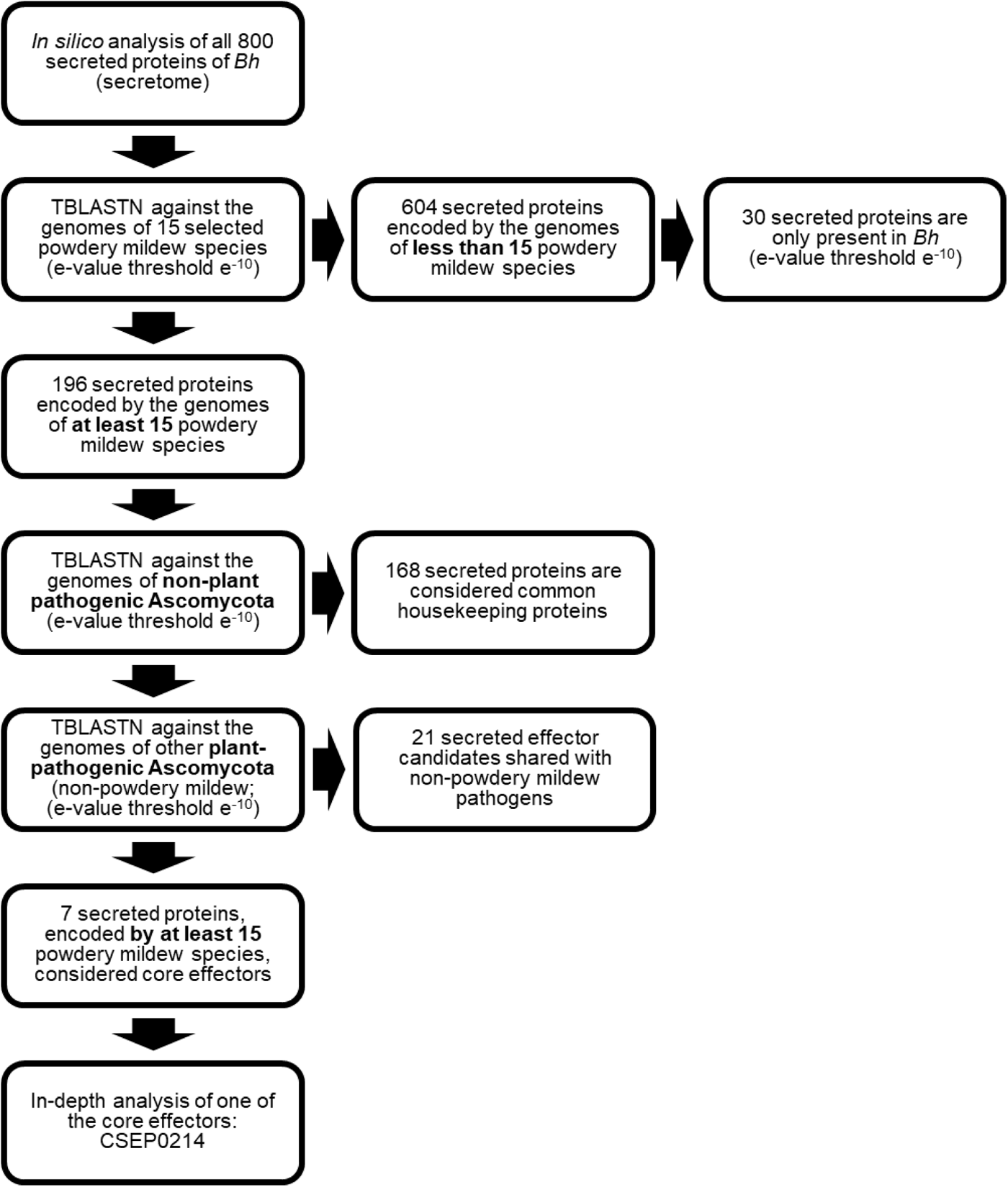
*In silico* analysis of the *B. hordei* secretome and identification of the powdery mildew-specific core effectome. All 800 proteins of *B. hordei* (*Bh*) that are predicted to be secreted were analysed *in silico* (see Supplementary File 1 and Supplementary Table 1) to obtain information on conserved domains and potential protein functions. TBLASTN searches were performed for these 800 *Bh* amino acid sequences against 17 genomes of 15 species from diverse phylogenetic clades of the *Erysiphaceae* (*B. graminis* f.sp. *triticale, B. graminis* f.sp. *tritici*, *Erysiphe alphitoides, E. necator*, *E. neolycopersici*, *E. pisi, Golovinomyces cichoracearum*, *G. magnicellulatus, G. orontii, Leveillula taurica*, *Parauncinula polyspora*, *Phyllactinia moricola*, *Pleochaeta shiraiana, Podosphaera leucotricha*, and *P. xanthii*). The 196 sequences found to be conserved in at least 15 out of the 16 genomes at an e-value threshold e^-10^ were searched by TBLASTN against three non-plant pathogenic ascomycetes (*Aspergillus* spp., *Penicillium* spp., *Saccharomyces* spp.), which identified 168 common potential fungal housekeeping proteins. The remaining 28 sequences were used for TBLASTN searches against a selection of plant-pathogenic ascomycetes (*Alternaria* spp., *Bipolaris* spp., *Botrytis* spp., *Claviceps* spp., *Colletotrichum* spp., *Curvularia* spp., *Drechslera* spp., *Drepanopeziza* spp., *Exserohilum* spp., *Fusarium* spp., *Gaeumannomyces* spp., *Magnaporthe* spp., *Monilinia* spp., *Pyrenophora* spp., *Pyricularia* spp., *Ramularia* spp., *Rhynchosporium* spp., *Sclerotinia* spp., *Septoria* spp., *Thielaviopsis* spp., *Venturia* spp., *Verticillium* spp., and *Zymoseptoria* spp.) to define the core effectome specific for the powdery mildew fungi. This revealed seven putative effector proteins that are exclusively present and highly conserved within the powdery mildew fungi (respective genes present in the genomes of at least 15 of the 16 (including *Bh*) analyzed powdery mildew species; see also Supplementary Table 2).

### CSEP0214 is a CFEM domain-containg protein and is highly expressed during fungal pathogenesis

The initial analysis described above, as well as TBLASTN searches in other powdery mildew fungal genomes, revealed that genes closely related to *Bh CSEP0214* are present in most of the tested powdery mildew fungal genomes, except in the early diverged and thus distantly related *P. polyspora*. We also failed to detect a CSEP0214 homologue in *Arachnopeziza araneosa*, a saprophyte that is the closest known relative of the powdery mildew fungi (Vaghefi et al., 2022) (Figure 2A). The amino acid alignment of CSEP0214 homologues revealed a high sequence conservation (49 to 80% identity) within the powdery mildew fungi (Figure 2B). Among four *Bh* isolates of different geographical origin (A6 (Denmark), DH14 (U.K.), K1 (Germany) and RACE1 (Japan)), the predicted amino acid sequence was homomorphic, indicating low within-species diversity of this candidate effector.

**Figure 2.**
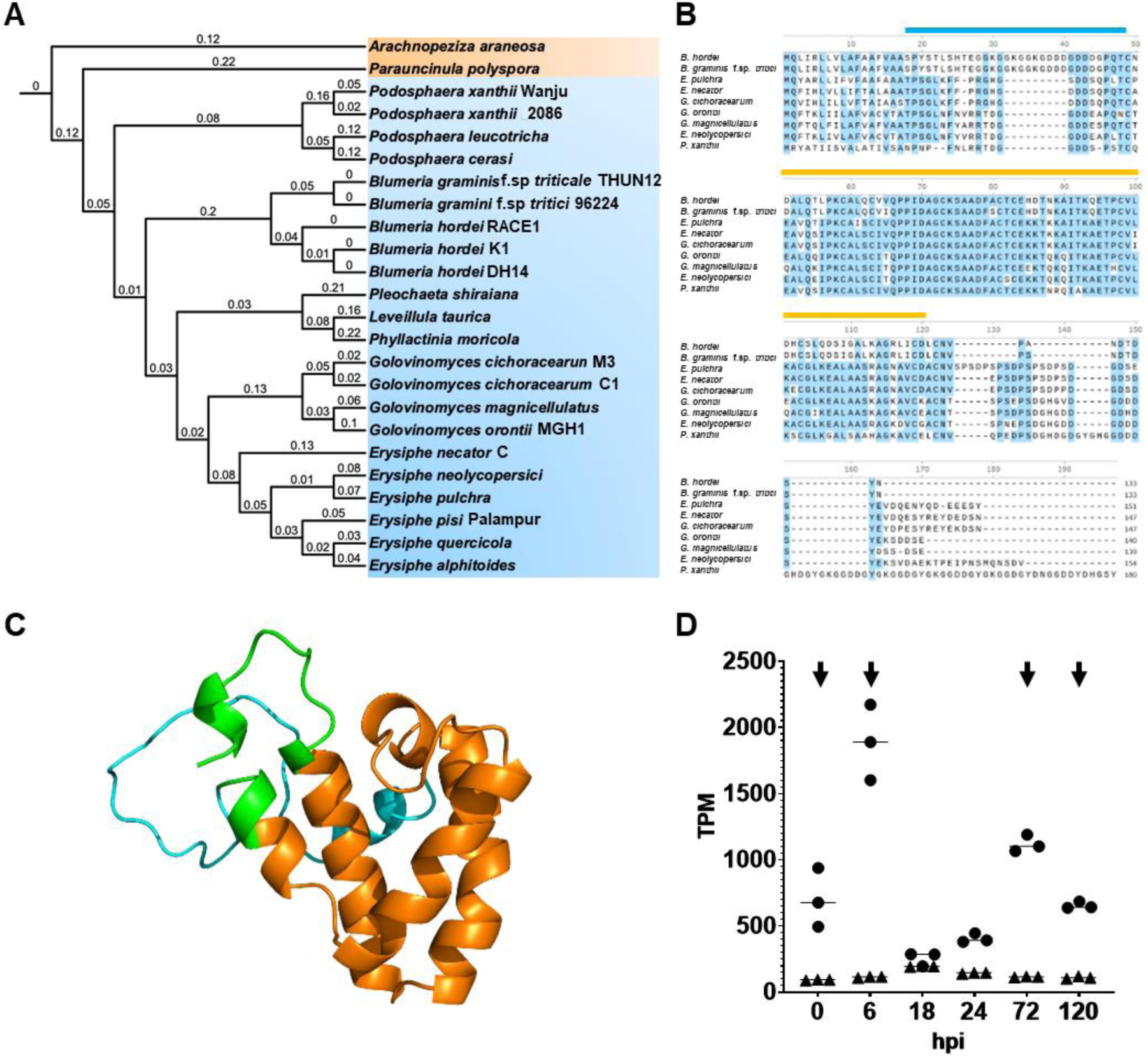
*In silico* analysis of CSEP0214. **A.** Phylogenetic tree (cladogram) of powdery mildew fungi based on the amino acid sequences of 129 homologous single copy proteins (Vaghefi et al., 2022). Numbers at branches refer to the average frequency of amino acid substitutions per site. Fungi with CSEP0214 homologues are shown in blue, fungi lacking CSEP0214 are depicted in red. **B.** Amino acid alignment of CSEP0214 homologues of nine powdery mildew species. Conserved amino acids (i.e., present in >55% of the sequences) are marked in blue. The blue bar above the alignment indicates the predicted disordered region and the orange bar indicates the predicted CFEM domain. **C.** Alphafold2-based prediction of the CSEP0214 three-dimensional structure; the predicted CFEM domain (orange) and the disordered region (light blue) as predicted by InterPro (https://www.ebi.ac.uk/interpro/search/sequence/) are highlighted. The protein regions outside these two domains are shown in green. **D.** Expression levels of *CSEP0214* (circles) in transcripts per million (TPM) based on three independent replicates of a time-course infection experiment of *Bh* isolate K1 on the susceptible barley cv. Margret compared to the average expression of transcripts of all 800 predicted secreted *Bh* proteins (triangles) at 0, 6, 18, 24, 72 and 120 hpi. Bars indicate medians. Arrows indicate time points where *CSEP0214* belongs to the top 30 expressed genes encoding secreted proteins. Whole transcriptome shotgun sequencing (RNA-Seq) data were retrieved from a previous study (Qian et al., 2023).

We predicted functional protein domains in CSEP0214 with InterPro (Blum et al., 2021) and detected an amino-terminal SP, comprising amino acids 1-17, followed by a predicted disordered region and a CFEM domain (Figure 2B). A CFEM domain is a fungus-specific protein domain with eight highly conserved cysteine residues and a proposed role in pathogenicity (Kulkarni et al., 2003). The greatest sequence conservation between the CSEP0214 homologues of powdery mildew fungi resides within this region (Figure 2B). Prediction of the three-dimensional structure of CSEP0214 by Phyre² and Alphafold2 enabled us to localize the position of the CFEM domain, the disordered region and undefined regions within the molecule (Figure 2C).

We used available (NCBI BioProject ID PRJNA835302; (Qian et al., 2023)) whole transcriptome shotgun sequencing (RNAseq) data to investigate the expression profile of *CSEP0214* during the infection of *Bh* on barley. In order to visualize the expression profile, we plotted the transcripts per million (TPM) of *CSEP0214* and the averaged TPM values of the transcripts of all proteins predicted to be secreted (800 proteins in total). This analysis revealed that the *CSEP0214* transcript accumulates at a level that is substantially higher than the average of the genes encoding secreted proteins. At 6 hours post inoculation (hpi), the difference is around 18-fold, whereas it almost decreases to the averaged TPM level for all secreted proteins at 18 hpi, only to increase again at 3 days after inoculation (Figure 2D). Overall, the *CSEP0214* transcript is among the 30 most abundant at both early (0 and 6 hpi) and late (72 and 120 hpi) time points during infection (Figure 2D).

### CSEP0214 interacts with the CORVET and HOPS complex component protein VPS18

To identify potential host target(s) of the mature full-length CSEP0214, we performed a classical Gal4-based yeast two-hybrid (Y2H) cDNA library screen and found a carboxy-terminal fragment of the barley RING domain protein, VPS18, as the most promising potential interaction partner (Supplementary Table 3). We recovered this carboxy-terminal fragment in eleven out of the 48 yeast colonies obtained in the Y2H screen. Cells of these eleven colonies activated the *HIS*, *ADE* and *lacZ* reporter genes and could grew on medium with up to 25 mM 3-amino-1,2,4-triazole (3-AT), indicating a strong protein-protein interaction (Supplementary Table 3). The shortest overlap encoded by the eleven prey clones covers VPS18 amino acids 842-992, which defines the interval required for the interaction with CSEP0214. This VPS18 carboxy-terminal 151-amino acid segment contains the complete RING domain of the protein, which is a core component of the HOPS and CORVET complexes. The CORVET complex is essential for the homotypic fusion between early endosomes or the heterotypic fusion between early endosomes and late endosomes/MVBs, while the HOPS complex is essential for the fusion of late endosomes and vacuoles. Thus, CORVET and HOPS serve as tethering factors for Rab5 and Rab7 GTPases, respectively (Seals et al., 2000; Peplowska et al., 2007; Balderhaar and Ungermann, 2013). For verification of the interaction between CSEP0214 and VPS18, we performed split-ubiquitin Y2H experiments with the 992-amino acid full-length (FL) version of VPS18. Combining VPS18, translationally fused to Cub-PLV at its carboxy-terminus, with NubG-CSEP0214, resulted in yeast growth on interaction-selective medium (Figure 3A), corroboarating the CSEP0214-VPS18 interaction. The split-ubiquitin Y2H system rather than the conventional Y2H system was chosen for this experiment as FL VPS18 did not express well in the latter.

**Figure 3.**
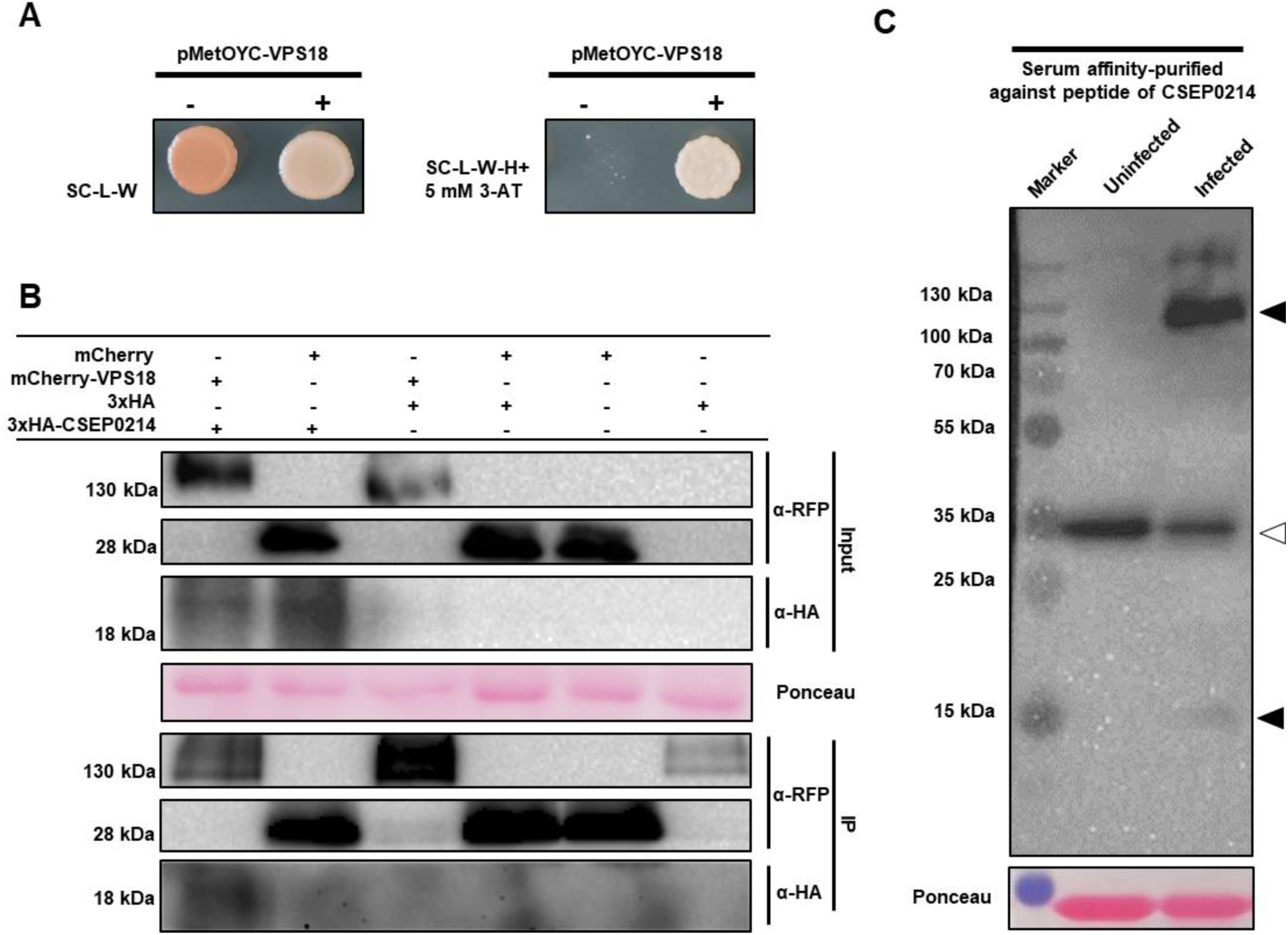
Interaction of CSEP0214 with VPS18. **A.** LexA-based split-ubiquitin Y2H assay of barley FL VPS18 expressed from vector pMetOYC (VPS18-Cub-PLV) and NubG-CSEP0214 (without SP) expressed from vector pNX32 (+) in the THY.AP4 strain. The empty pNX32 vector was used as negative control (-). Yeast growth on medium that is selective for the presence of the plasmids (SC-L-W; growth control) is shown on the left, and yeast growth on interaction-selective medium (SC-L-W-H+5 mM 3AT) is shown on the right, after 3 days of growth. Experiments were performed three times with similar results. **B.** Fusion proteins mCherry-VPS18 and 3xHA-CSEP0214, or the mCherry and 3xHA tags alone were co-expressed in different combinations via *A. tumefaciens*-mediated transient gene expression in leaves of *N. benthamiana* plants. Proteins were extracted for Co-IP. Protein samples of the IP and the total protein extract (Input) were used for SDS-PAGE and immunoblot analysis. The membranes were probed with *α-*RFP and *α-*HA antibodies, respectively. On the left, molecular masses of marker proteins are given. Ponceau staining was used to demonstrate equal amounts of total proteins on the blots. **C.** Immunoblot detection of CSEP0214 in total protein extracts of *Bh*-infected (5 dpi with virulent isolate K1) and uninfected barley cv. Lottie leaves. Affinity-purified antiserum raised against CSEP0214 (see Materials and Methods for details), was used. The white arrowhead marks an unspecific band present in samples from both non-infected and infected leaves, black arrowheads mark bands that are detectable solely in samples from infected leaves. On the left, molecular masses of marker proteins are given. Ponceau staining was used to demonstrate equal amounts of total proteins on the blots.

We further performed co-immunoprecipitation (Co-IP) experiments to validate our yeast-based results. To this end, we transiently co-expressed fluorophore (mCherry)-tagged VPS18 with triple hemagglutinin (3xHA)-tagged CSEP0214 (lacking its SP) or the respective empty vectors (as negative controls), in different combinations in leaves of *Nicotiana benthamiana* and used red fluorescent protein (RFP) trap beads to purify potential protein complexes from the respective plant lysates. We only recovered 3xHA-CSEP0214 when co-expressed with mCherry-VPS18 (Figure 3B). Taken together, both assays (split-ubiquitin Y2H and Co-IP) provided strong evidence for an interaction between CSEP0214 and full-length barley VPS18.

We next tested if the interaction between CSEP0214 and VPS18 is specific. Therefore, we performed a classical Y2H experiment with the 85-amino acid RING domain fragment of VPS18, originally recovered in the Y2H screen (see above), against a test set of ten randomly selected *Bh* CSEPs. As judged from yeast growth on interaction-selective medium, and in contrast to CSEP0214, none of these CSEPs were able to interact with the VPS18 RING domain (Supplementary Figure 1A). To demonstrate that a lack of interaction in the Y2H assay was not due to failed expression of the CSEPs, we performed immunoblot analysis and found that eight of the CSEPs were detectably expressed, albeit at different levels (Supplementary Figure 1B). To explore whether CSEP0214 interacts specifically with the RING domain of VPS18, we tested the RING domains of barley PEX2 and PEX12 (see sequence alignment in Supplementary Figure 1C), two barley peroxins involved in mediating peroxisomal protein import (Kao et al., 2016). Neither of these showed signs of interaction in the classical Y2H assay (Supplementary Figure 1D).

We raised a polyclonal peptide antiserum against CSEP0214 (Supplementary Table 4) and affinity-purified it against the peptide used for immunization (for details, see Materials and Methods). This purified antiserum was tested using 6x-His-CSEP0214 protein inducibly expressed in *Escherichia coli*, and we detected a band corresponding to the molecular mass of CSEP0214 (∼12 kDa) in bacterial lysates (Supplementary Figure 2). The affinity-purified antiserum was subsequently used in immunoblots to analyze if we can detect CSEP0214 in protein samples from *Bh*-infected barley leaves. Besides an unspecific binding product that was present in both non-inoculated and *Bh*-inoculated leaf samples, we detected a weak band corresponding to the calculated molecular mass of CSEP0214 (∼12 kDa) specifically in inoculated leaf samples. We further observed stronger bands of high molecular mass (at ∼130 kDa). These were likewise only present in samples from *Bh*-inoculated leaves and could correspond to a complex composed of the 992-amino acid VPS18 (∼109 kDa) and CSEP0214 (Figure 3C).

Having found that CSEP0214 and VPS18 might form a stable complex *in planta*, we wanted to analyze the strength of this interaction in more detail. Therefore, we performed an immunoblot experiment using combinations of mCherry-VPS18 and 3xHA-CSEP0214 as well as empty vector controls expressed in leaves of *N. benthamiana*. The proteins were extracted using either a standard extraction buffer or a denaturation buffer containing high concentrations of urea and thiourea (“urea buffer”), and separated by sodium dodecyl sulfate-polyacrylamide gel electrophoresis (SDS-PAGE). On an anti-HA immunoblot, we detected an effector-specific band of ∼18 kDa. However, in the sample extracted with the mild buffer, and not with the denaturation buffer, we also detected a band of ∼140 kDa using the anti-HA antibody (Supplementary Figure 3). This outcome is indicative of a heat- and SDS-resistant VPS18-CSEP0214 complex that can only be dissolved by strong denaturing agents.

### The C-terminal portion of the CFEM domain in CSEP0214 and the RING domain of VPS18 are essential for the interaction

To narrow down the potential interaction site within CSEP0214, we created seven truncated versions of the effector protein (Figure 4A) and analyzed these for interaction with FL VPS18 in split-ubiquitin Y2H experiments. Here we observed that FL CSEP0214 and surprisingly all truncated versions as well showed interaction with FL VPS18. These truncated variants included at least three essentially non-overlapping CSEP0214 domains (CSEP0214^1-34^, CSEP0214^31-71^ and CSEP0214^85-107^) (Figure 4B). When we used the classical Y2H system to test for interaction between VPS18 RING domain and these CSEP0214 truncations, we only saw interaction when CSEP0214 amino acids 85-107 were included but not with the fragments located N-terminally hereof. Thus, CSEP0214^1-34^, CSEP0214^1-85^, CSEP0214^31-71^ and CSEP0214^1-71^ failed to interact with the VPS18 RING domain (Supplementary Figure 4A). To validate this result, we confirmed expression of the truncated CSEPs in the yeast strains by immunoblot-based detection of the Gal4-binding domain of the various constructs (Supplementary Figure 4B). Altogether, the results suggest that the C-terminal region of the CFEM domain in CSEP0214 (between amino acids 85 and 107) is essential for the interaction with the VPS18 RING domain. However, the remaining part of CSEP0214 is likely involved in interaction with regions in VPS18 other than the RING domain. Returning to the split-ubiquitin Y2H system, we indeed saw interaction between VPS18 ΔRING (lacking the C-terminal region with the RING domain) and FL CSEP0214 (Figure 4C and D, Supplementary Figure 5). All-in-all, the studies suggest a complex interaction between the two proteins, involving at least three CSEP0214 and two VPS18 domains.

**Figure 4.**
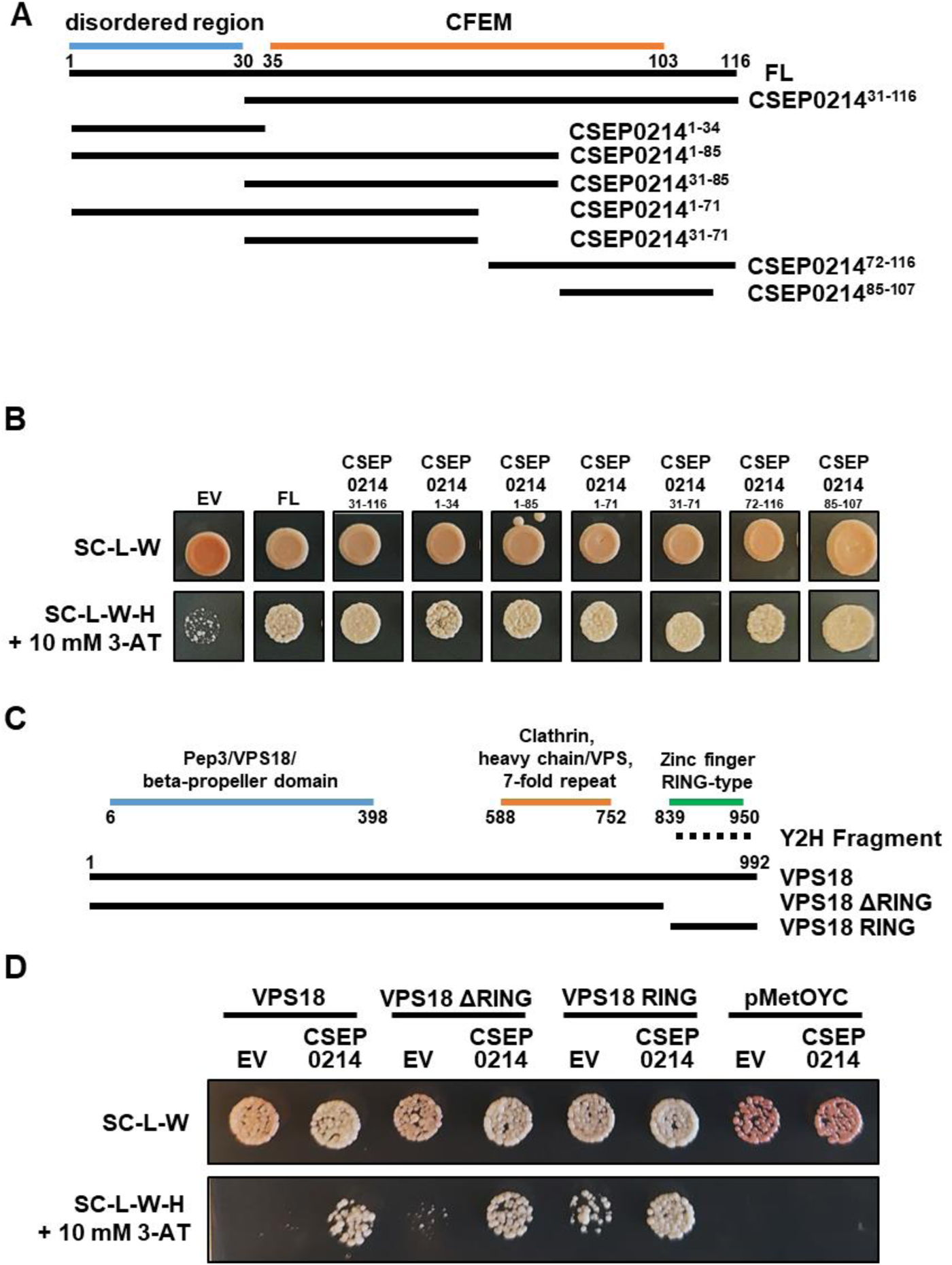
The C-terminal portion of the CFEM domain in CSEP0214 interacts with the VPS18 RING domain. **A.** Schematic representation of different truncated versions of CSEP0214 and indication of predicted domains. FL: full-length (without SP, 116 amino acids). The indicated numbering of the amino acids refers to this 116-aa version. **B.** LexA-based split-ubiquitin Y2H assay of barley FL VPS18 expressed from pMetOYC (VPS18-Cub-PLV) and truncated versions of CSEP0214 expressed from pNX32 (NubG-CSEP0214) in the THY.AP4 strain. The empty pNX32 (NubG-GWY) vector served as a negative control (EV). Aliquots of yeast liquid cultures were dropped on either SC-L-W medium (growth control) or SC-L-W-H + 10 mM 3-AT medium (interaction-selective) (**B**, **D**). **C.** Schematic representation of VPS18 and two truncated versions (VPS18 ΔRING and VPS18 RING) and location of predicted domains. The numbering indicates the amino acid position of predicted domains. Dashed line depicts the fragment identified in the Y2H cDNA screen. **D.** LexA-based split-ubiquitin Y2H assay with FL and truncated versions of VPS18 expressed from pMetOYC (VPS18-Cub-PLV) and FL CSEP0214 expressed from pNX32 (NubG-CSEP0214). Images were taken after 3 days of growth.

### Expression of CSEP0214 in barley leaf epidermal cells or knockdown of its target perturbs the endomembrane trafficking pathway

As VPS18 is known to be a core component of the CORVET and HOPS complexes involved in the vacuolar transport pathway (van der Kant et al., 2015), we wanted to examine if CSEP0214 affects this route via interaction with VPS18. Thus, we transiently expressed CSEP0214 lacking its SP (all following experiments were performed with this version of the effector) and the vacuolar marker (SP)-RFP-AFVY, where the four-amino acid vacuolar targeting signal of the vacuolar lumen protein, phaseolin (Hunter et al., 2007), was added to the C-terminus of red fluorescent protein, in barley leaf epidermal cells. We observed that expression of CSEP0214 caused a significant fraction of this vacuolar marker protein to be retained in an endoplasmic reticulum (ER)-like reticulate structure (Figure 5A). The ER identity of this structure was confirmed by co-localization with a fluorescent luminal ER marker, (SP)-mYFP-HDEL (i.e., monomeric yellow fluorescent protein (YFP) with the carboxy-terminal HDEL ER retention signal; (Irons et al., 2003)) in 47 out of 50 cells studied, as compared to three out of 50 in the empty vector control (Figure 5A).

**Figure 5.**
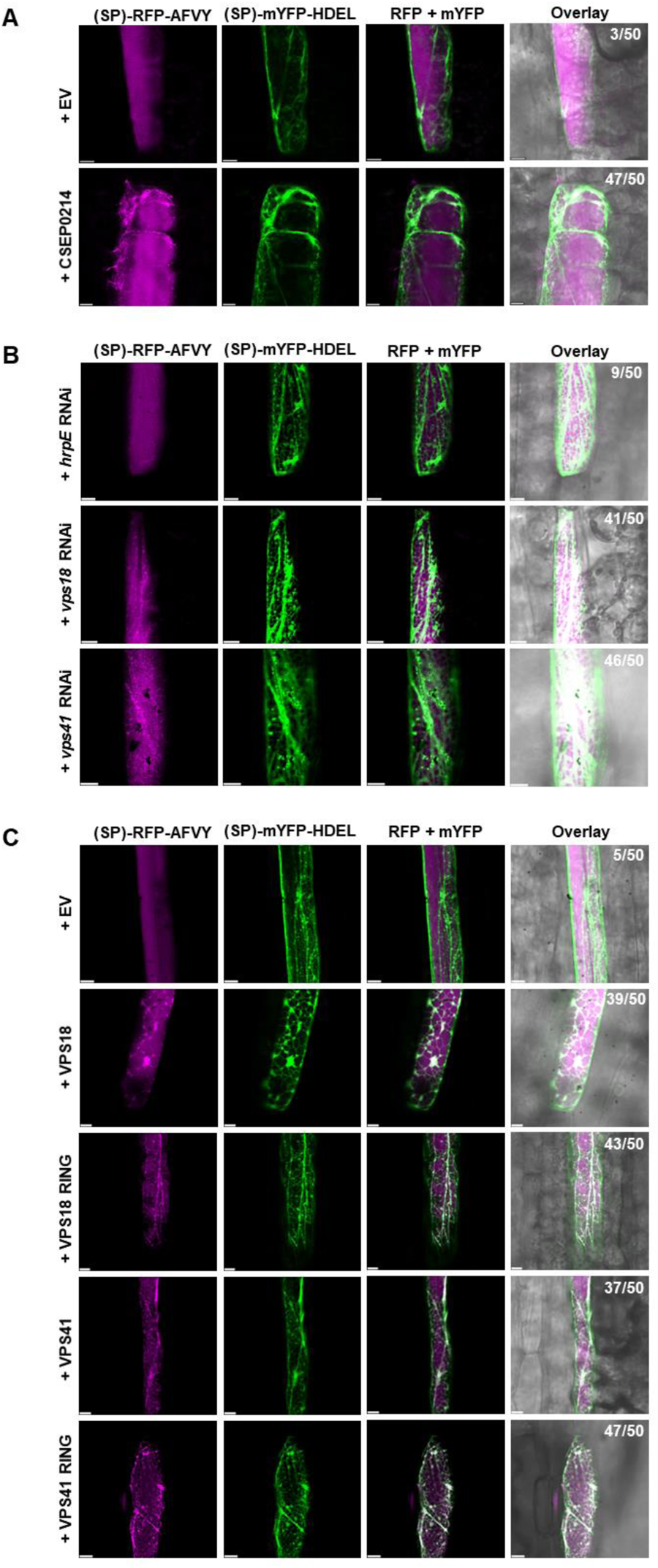
Expression of CSEP0214 in barley leaf epidermal cells or knockdown of the target blocks the transport of the vacuolar marker protein. Subcellular localization of the vacuolar marker (SP)-RFP-AFVY and the ER marker (SP)-mYFP-HDEL upon co-expression with either (**A**) an empty vector (EV; pUbi::GW) or CSEP0214 (pUbi::CSEP0214), (**B**) the *hrpE* control RNAi construct (pIPKTA30N::*hrpE*) or knockdown constructs of *VPS18* (pIPKTA30N::*VPS18*) and *VPS41* (pIPKTA30N::*VPS41*), or (**C**) an empty vector (EV; pUbi::GW) or expression of VPS18 (pUbi::VPS18), VPS18 RING domain (pUbi::VPS18 RING), VPS41 (pUbi::VPS41) or VPS41 RING domain (pUbi::VPS41 RING), in barley (cv. Golden Promise) by particle bombardment of leaf epidermal cells. Numbers in the top right corner of the overlay panels indicate the occurrence of co-localization with the ER marker in 50 inspected cells. Scale bar, 10 µm.

To support that CSEP0214 acts via VPS18, we used an RNA interference (RNAi) approach targeting *VPS18* and *VPS41*. In these experiments, we bombarded barley leaf epidermal cells with RNAi constructs, designed for the knockdown of *VPS18* and *VPS41*, along with the vacuolar marker (SP)-RFP-AFVY. The *hrpE* gene from the type-III secretion system of the bacterial phytopathogen *Pseudomonas syringae* was used as a non-plant RNAi control. We observed specifically that RNAi-based knockdown of *VPS18* or *VPS41* caused retention of the fluorescent vacuolar marker in reticulate structures that co-localized with the ER marker, (SP)-mYFP-HDEL, similar to the effect seen with the overexpression of CSEP0214 (Figure 5B). In order to further confirm this observation, we wanted to test if putative dominant-negative variants of VPS18 and VPS41 affected this route. Thus, we transiently expressed the RING domains, and the full-length versions of these proteins as controls, along with the vacuolar marker, (SP)-RFP-AFVY, in barley leaf epidermal cells. In all four cases, this led to ER localization of the vacuolar marker (Figure 5C). Expression of FL VPS18 or VPS41 also blocked (SP)-RFP-AFVY in the ER, indicating that correct stoichiometry of the components in the CORVET and/or HOPS complexes is essential for proper function in endomembrane trafficking (Figure 5C). Moreover, RNAi of the CORVET-specific *VPS8* showed a similar block of (SP)-RFP-AFVY in the ER (Supplementary Figure 6A).

In addition, expression of CSEP0214 caused a partial overlap between the fluorescent Golgi ST-YFP marker (a fusion of the 52-amino acid signal anchor of a rat sialyltransferase fused to YFP (Brandizzi et al., 2002)) and the ER marker in 32 out of 50 cells studied, as compared to 0 out of 50 in case of the empty vector control (Supplementary Figure 7A). Furthermore, CSEP0214 expression blocked the otherwise secreted protein marker, (SP)-mCherry, and caused it to accumulate in intracellular aggregates and an ER-like structure (Supplementary Figure 7B).

### Expression of CSEP0214 in barley leaf epidermal cells or knockdown of its target blocks the transport and accumulation of the papilla marker protein, ROR2

In order to explore whether CSEP0214 affects a well-known immunity-associated endomembrane trafficking pathway, the transport of marker proteins upon *Bh* challenge to the papilla was examined. We co-expressed the established papilla marker, mCherry-ROR2 (Bhat et al., 2005), with CSEP0214 in barley leaf epidermal cells, followed by *Bh* inoculation. In the empty vector control, mCherry-ROR2 localized at the plasma membrane and in papillae formed at *Bh* attack sites (Figure 6A). This strong mCherry-ROR2 labelling of extracellular papillae agrees with previous observations that this membrane protein and its orthologue, Arabidopsis PEN1, are markers for extracellular vesicles (Meyer et al., 2009; Böhlenius et al., 2010; Nielsen et al., 2012). However, when co-expressed with CSEP0214, mCherry-ROR2 did not localize at papillae, but instead appeared in discrete structures, accumulating near the fungal attack site and to some extent in ER-like structures, in 46 out of 50 cells studied, as compared to 8 out of 50 in the empty vector control (Figure 6A). In the absence of *Bh* challenge, CSEP0214 expression led to localization of ROR2 in a reticulate structure overlapping with the ER marker (Supplementary Figure 7C). The observed effect of the CSEP0214-induced block of ROR2 transport papillae could also be seen upon RNAi of *VPS18* or *VPS41* (Figure 6B), or RNAi of *VPS8* (Supplementary Figure 6B). Co-expression of either VPS18, VPS18-RING, VPS41 or VPS41-RING also blocked ROR2 localization to papillae, and mCherry-ROR2 was observed in an ER-like structure rather than in the papillae, indicating that these proteins are important for this cellular trafficking pathway (Figure 6C).

**Figure 6.**
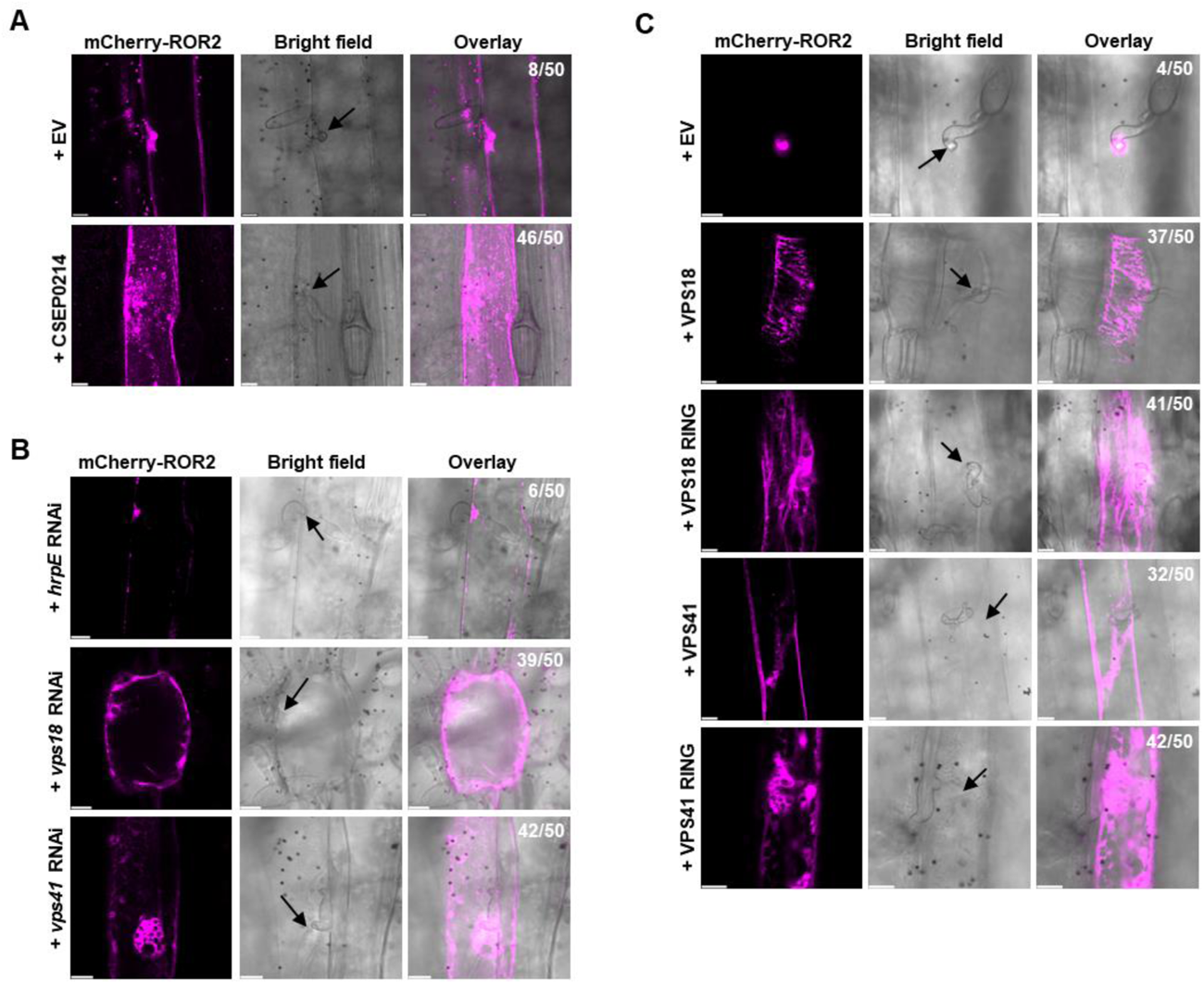
Expression of CSEP0214 in barley leaf epidermal cells or knockdown of the target blocks the transport and accumulation of the papilla marker protein, ROR2. Subcellular localization of the papilla marker mCherry-ROR2 upon co-expression with either (**A**) an empty vector (EV; pUbi::GW) or CSEP0214 (pUbi::CSEP0214), (**B**) the *hrpE* control RNAi construct (pIPKTA30N::*hrpE*) or knockdown constructs of *VPS18* (pIPKTA30N::*VPS18*) and *VPS41* (pIPKTA30N::*VPS41*), or (**C**) an empty vector (EV; pUbi::GW) or expression of VPS18 (pUbi::VPS18), VPS18 RING domain (pUbi::VPS18 RING), VPS41 (pUbi::VPS41) or VPS41 RING domain (pUbi::VPS41 RING), in barley (cv. Golden Promise) by particle bombardment of leaf epidermal cells. Arrows indicate fungal attack sites. Numbers in the top right corner of the overlay panels indicate the occurrence of unusual marker localization in 50 inspected cells. Scale bar, 10 µm.

### CSEP0214 mediates susceptibility by blocking immune receptor-induced hypersensitive response and the encasement of fungal infection structures

Previously, MON1, involved in the activation of the late endosome GTPase Rab7, has been found to be important for encasement formation and the hypersensitive response (HR) in response to powdery mildew fungi in barley and *Arabidopsis thaliana* (Liao et al., 2023). Since we provide data above indicating that CSEP0214 affects HOPS, the tethering complex of Rab7, we analyzed whether CSEP0214 hampered these immune responses as well. To test for the occurrence of HR, we used the barley lines P01 and P02, which are immune to *Bh* isolates C15 and A6, respectively. The isolate-specific immunity in these lines is mediated by the allelic *Mla1* (P01) and *Mla3* (P02) resistance genes, encoding coiled-coil, nucleotide-binding, leucine-rich repeat (CNL)-type resistance proteins (Seeholzer et al., 2010; Kølster et al., 1986). We used the bacterial *Pseudomonas fluorescence* strain EtHAn, engineered to express a *P. syringae* type 3-secretion system, and the pEDV6 vector (Thomas et al., 2009; Fabro et al., 2011), to introduce either β-glucuronidase (control) (Li et al., 2024), or CSEP0214 into leaf cells of barley lines P01 or P02 by infiltrating the bacteria into the leaf apoplastic space. The leaves were subsequently inoculated with the avirulent *Bh* isolates, C15 and A6, respectively. Samples were collected at four days after infiltration/inoculation and stained with either Coomassie blue (Figure 7A, C, E, G) to visualize fungal hyphae, or with trypan blue (Figure 7B, D, F, H) to detect host cell death. Pathogenic growth of the fungus was significantly increased, quantified as secondary hyphal growth and conidiophore production, after delivery of CSEP0214 (Figure 7I, J). In addition, CSEP0214 suppressed the HR-like cell death in these otherwise resistant barley lines (Figure 7K).

**Figure 7.**
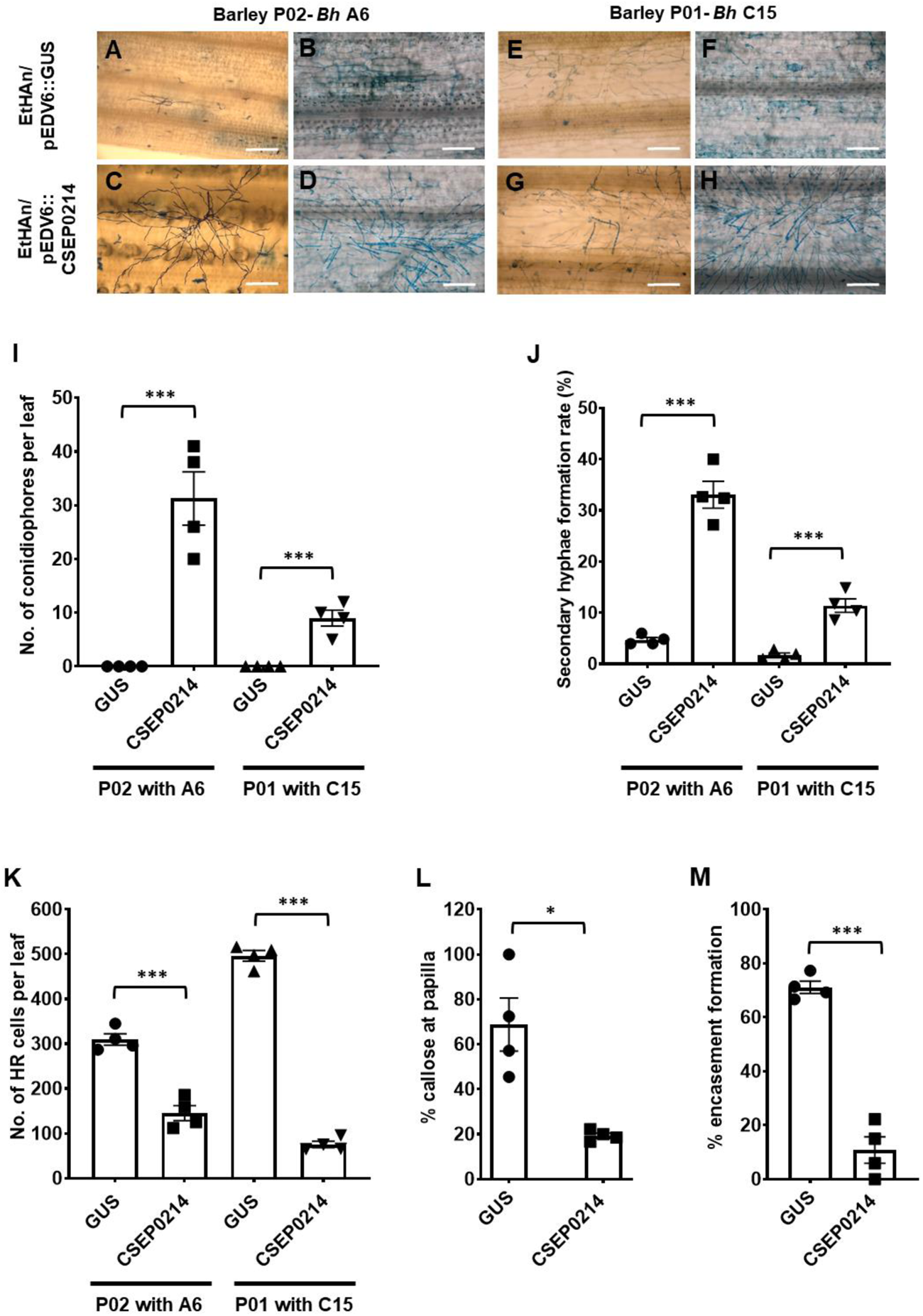
Expression of CSEP0214 inhibits CNL-mediated resistance, as well as papilla and encasement formation in response to *Bh*. A-K. *Mla3* (in P02) and *Mla1* (in P01)-mediated resistance to *Bh* in barley is broken by expression of CSEP0214. The EtHAn strain of *P. fluorescence* was used to introduce either GUS, as a control, or CSEP0214 into leaves of barley lines P02 or P01. The leaves were subsequently inoculated with the avirulent *Bh* isolates, A6 and C15, respectively. Samples were collected at 4 dpi and stained with either Coomassie blue (**A, C, E, G**) to visualize the fungal hyphae, or with trypan blue (**B, D, F, H**) to stain for cell death. Pathogenic success of the fungus was quantified as the number of conidiophores per leaf of equal size (**I**) and the percentage of germinated spores forming hyphae (**J**). Cell death was measured as the number of cells showing HR per leaf of equal size, after staining with trypan blue (**K**), 4 days post inoculation. **L.** Effect of CSEP0214 expression on callose deposition at papillae. Leaf epidermal cells of seven-day-old barley (cv. Golden Promise) plants were transformed by particle bombardment with a construct expressing GUS along with either empty vector (EV; pUbi::GW) or expression of CSEP0214 (pUbi::CSEP0214). After one day, the leaves were inoculated with *Bh* (isolate C15). Percentage of GUS-positive cells with callose in the papilla in barley leaf epidermal cells was scored at 24 hpi, by GUS staining, followed by aniline blue staining and observation with UV-fluorescence microscopy. **M.** Effect of CSEP0214 expression on encasement of *Bh* (isolate C15) haustoria. Leaf epidermal cells of seven-day-old barley (cv. Golden Promise) plants were transformed as in ‘**L**’, treated with tetraconazole (100 µM) and inoculated with *Bh* (isolate C15) 2 h later. After 4 days, the percentage of GUS cells with callose-encased haustoria was scored, by observation with UV-fluorescence microscopy after aniline blue staining. n = 4. Error bars, SE. *, P<0.05; ***, P<0.001 assessed by Student’s t-tests.

We next analyzed the extent of callose deposition in papillae upon transient expression of CSEP0214. Either empty vector or a construct for expression of untagged CSEP0214 were introduced into barley leaf epidermal cells by particle bombardment along with a construct expressing β-glucuronidase to allow detection of transformed cells. In this assay, we observed a significant reduction in callose deposition in papillae following expression of CSEP0214 in the leaf cells (Figure 7L). *Bh* normally does not induce encasement formation in barley. However, this immune-related structure is formed when the plants are sprayed with the fungicide, tetraconazole (Maffi et al., 1995; Liao et al., 2023). The formation of tetraconazole-induced encasements was significantly reduced following CSEP0214 expression (Figure 7M). To analyse whether the reduced callose response affected the fungal host cell entry rate, we introduced constructs for expression of CSEP0214, VPS18 and the VPS18 RING domain by particle bombardment. None of the three constructs caused a statistically significant difference from the empty vector control regarding the rate of haustorium formation (Supplementary Figure 8).

### Expression of CSEP0214 inhibits interaction of VPS18 with VPS16

In order to investigate further how CSEP0214 inhibits the function of VPS18, we wanted to analyze if CSEP0214 inhibits the interaction between VPS18 and VPS16, another component of the CORVET and HOPS complex. These two proteins have earlier been shown to associate (Graham et al., 2013; van der Kant et al., 2015; Shvarev et al., 2022). We performed split-ubiquitin Y2H experiments with VPS18 (FL), fused to Cub-PLV (pMetOYC), against VPS16 (FL)-NubG (pXN22). When co-expressed with empty vector (pAG426-GPD-GW) (Alberti et al., 2007), we observed yeast growth on selective medium indicating interaction between VPS18 and VPS16 (Figure 8A). However, when CSEP0214 was co-expressed from pAG426-GPD-CSEP0214 along with VPS18-VPS16, yeast growth was reduced, indicating a diminshed interaction between VPS18 and VPS16 (Figure 8A).

**Figure 8.**
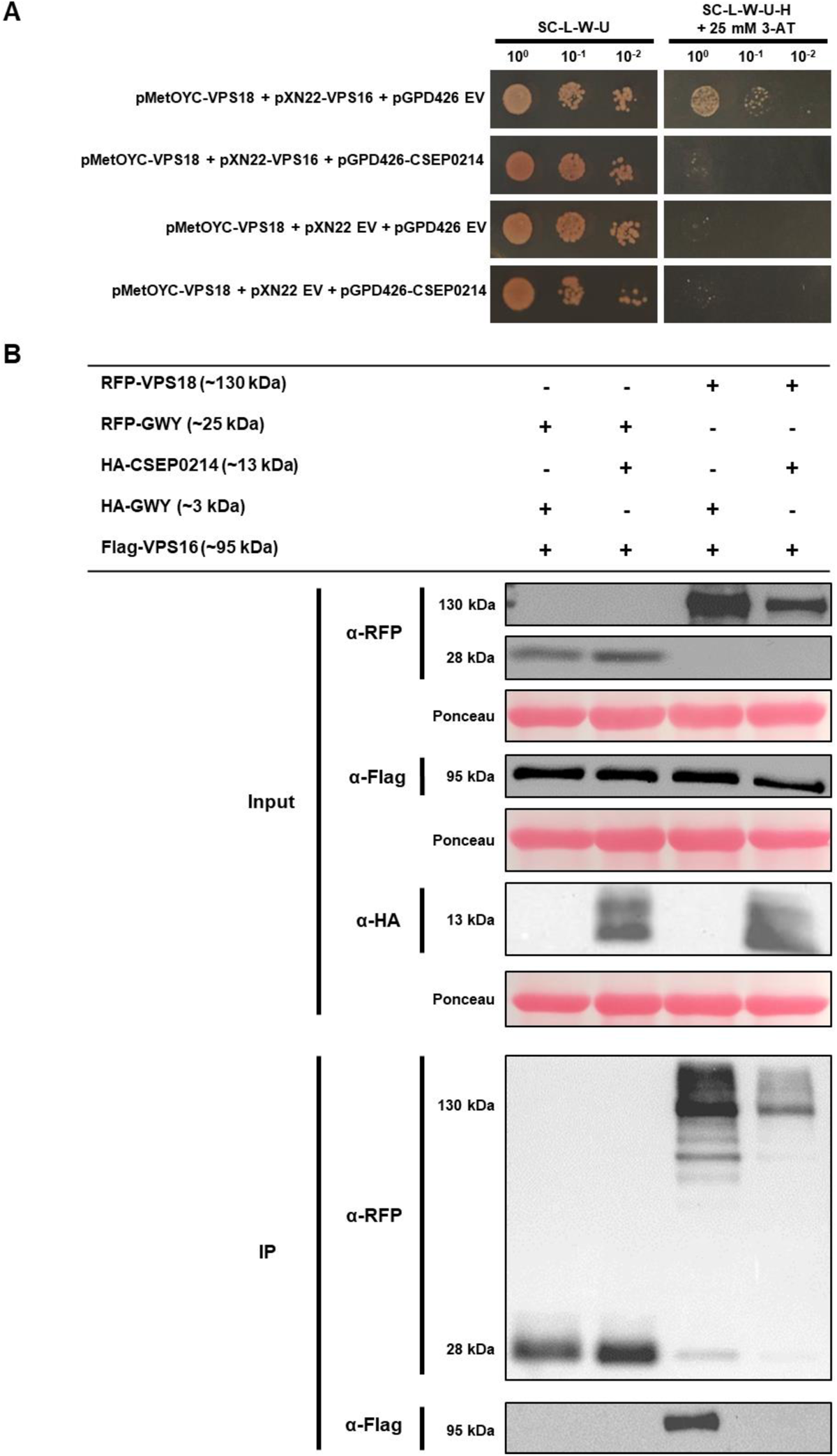
Expression of CSEP0214 inhibits interaction of VPS18 with VPS16. **A.** Three-way split-ubiquitin Y2H of full-length barley VPS18 expressed from pMetOYC (VPS18-Cub-PLV) and full-length barley VPS16 expressed from pXN22 (VPS16-NubG), in the presence or absence of pAG426-GPD-GW empty vector (EV), or pAG426-GPD-CSEP0214 in the THY.AP4 strain. pMetOYC-VPS18 + EV pXN22 + EV pAG426-GPD-GW, and pMetOYC-VPS18 + EV pXN22 + pAG426-GPD-CSEP0214 served as negative controls. Aliquots of yeast liquid cultures were dropped in various dilutions on either SC-L-W-U medium (growth control) or SC-L-W-U-H + 25 mM 3-AT medium (interaction-selective). Images were taken after 2 days of growth. **B.** Fusion proteins mRFP-VPS18, HA-CSEP0214 and Flag-VPS16 or the mRFP and HA tags alone, were co-expressed in different combinations via *A. tumefaciens*-mediated transient gene expression in leaves of *N. benthamiana* plants. Proteins were extracted for Co-IP (co-immuno-precipitation (IP). Protein samples of the IP and the total protein extract (Input) were used for SDS-PAGE and immunoblot analysis. The membranes were probed with *α-*RFP, *α*-Flag and *α-*HA antibodies, respectively. On the left, molecular masses of marker proteins are indicated. Ponceau staining was used to demonstrate equal amount of total proteins on the blots.

To validate this interference *in-planta*, we co-expressed either RFP or RFP-VPS18 with Flag-VPS16 in *N. benthamiana* leaves, in the presence or absence of HA-tagged CSEP0214 (Figure 8B). We used RFP trap beads to co-purify the interacting partners from the cell lysate. We detected Flag-VPS16 in the IP blot with RFP-VPS18, but only when HA-CSEP0214 was absent (Figure 8B). These Y2H and Co-IP experiments indicate that CSEP0214 blocks the interaction between VPS18 and VPS16.

## Discussion

In this study we identified the CORVET and HOPS complex component VPS18 as a cellular target of the *Bh* core effector protein CSEP0214 (BLGH_02334). The genomes of powdery mildew fungi encode predicted effector arsenals that range from fewer than 100 putative effector genes in *P. polyspora* (Frantzeskakis et al., 2019) to more than 600 in the cereal-infecting pathogens of the genus *Blumeria* (Frantzeskakis et al., 2018; Müller et al., 2018), reviewed in (Barsoum et al., 2019). Here, we explored conservation of effector genes across the entire powdery mildew fungal lineage and identified seven genes that encode highly conserved effector proteins present in at least fifteen out of the sixteen powdery mildew fungal species analyzed (Figure 1 and Supplementary Table 2). These effectors form the powdery mildew-specific core effectome expected to fulfil essential functions during host colonization and thus contribute to pathogenic success. As a result of this investigation, we also provide the scientific community with an in-depth analysis of the secretome of *Bh*, which updates and extends the results of a former study (Supplementary Table 1) (Pedersen et al., 2012).

In view of the highly similar lifestyle and mode of pathogenicity of powdery mildew fungi and given the comparatively high number of effector genes in these species, the limited size of the identified core effectome, specific for this pathogen clade, is surprising. We cannot exclude that we underestimate the extent of the powdery mildew-specific core effectome due to the in part limited quality of the currently available powdery mildew genome assemblies. Nonetheless, it was already suggested that pathogenicity of powdery mildew fungi rests on a narrow core effectome that is augmented by a variable number of species/lineage-specific effectors required for the colonization of the various host plants (Liang et al., 2018).

We selected CSEP0214 (BLGH_02334), one of the seven powdery mildew core effectors, for detailed functional analysis. This effector was previously recognized as conserved between *B. graminis,* the Arabidopsis powdery mildew pathogen, *Golovinomyces orontii* and the pea powdery mildew *Erysiphe pisi* (Schmidt et al., 2014). Using an Y2H screen against a barley prey cDNA library, we identified VPS18 as a host protein that interacts with this effector. The CSEP0214-VPS18 interaction was seen in two different yeast systems and in biochemical Co-IP experiments (Figure 3A and B, Supplementary Figure 2A), and we provide evidence that it is mediated by a number of molecular contacts along the two proteins (Figure 4 and Supplementary Figure 4). The interaction is further supported by an SDS- and heat-resistant high-molecular-weight band in protein extract from infected barley leaves, detected with a CSEP0214-specific antiserum (Figure 3C). The molecular mass of this band is consistent with the estimated mass of a VPS18-CSEP0214 complex, which is hard to separate due to intricate molecular interactions, for instance involving the C-terminal RING domain of VPS18 and part of the CFEM domain of CSEP0214. Some CFEM domain-containing effectors are critical for fungal virulence (Bai et al., 2022; Wang et al., 2022; Zhu et al., 2017; Zuo et al., 2022; Shang et al., 2024). In the grass pathogen *Fusarium graminearum*, for example, multiple CFEM effectors are essential for full virulence on maize as they negatively regulate the activity of a host cell wall-associated kinase (Zuo et al., 2022).

Membrane trafficking is a key process in plant-microbe interactions that is heavily targeted by effectors (Inada and Ueda, 2014; Bhandari and Brandizzi, 2024). It is required for the internalization of cell surface receptors (Beck et al., 2012), the secretion of antimicrobial cargo (Baena et al., 2022; Kim et al., 2014) and cell wall components (Sinclair et al., 2018), and possibly the accommodation of infection structures of certain pathogens (Abubakar et al., 2023; Berkey et al., 2017). VPS18 is a highly-conserved core component of the CORVET and HOPS complexes that regulate endocytic membrane trafficking towards the plant vacuole, and the protein interacts with other HOPS and CORVET components, e.g. VPS16, VPS41, and VPS33, via its RING domain (Takemoto et al., 2018; Hou et al., 2021; Brillada et al., 2018). While oomycete pathogens employ effectors to redirect this endomembrane traffic towards their haustoria to generate the extrahaustorial membrane and bring resources to sustain them (Gu et al., 2017; Yuen et al., 2023; Jeon and Segonzac, 2023; Bozkurt et al., 2015; Petre et al., 2021), we observe that the barley powdery mildew fungus inhibits this pathway. The high expression level of CSEP0214 in the first six hours after inoculation indicates that this inhibition is essential from the early phase of infection. The high amino acid conservation of this effector between powdery mildew fungi, which aligns with the highly conserved target, VPS18, suggests that this inhibition is essential across powdery mildew fungal species. Our study of the truncations of the CSEP0214 also indicate that this interaction is extremely strong, and spans the entire protein (Figure 4). Thus, the identification of the CSEP0214-VPS18 interaction extends the growing list of plant endomembrane trafficking components that are targeted by powdery mildew effectors. For example, the *Bh* effector candidate BEC4 binds a barley ADP ribosylation factor-GTPase activating protein (ARF-GAP), thereby possibly affecting defence-associated vesicle trafficking in the host (Schmidt et al., 2014). Additionally, the *Bh* effector CSEP0162 associates with barley MON1, a component that is important for fusion of MVBs to their target membranes (Liao et al., 2023).

Here, we investigated membrane trafficking to the vacuole in relation to the effect of CSEP0214 on VPS18. Overall, we found that both expression of CSEP0214 and silencing of *VPS18* and other CORVET and HOPS components in barley cells impeded trafficking of the vacuolar marker protein, (SP)-RFP-AFVY (Figure 5), the secreted protein, (SP)-mCherry (Supplementary Figure 7C), the Golgi marker protein, ST-mYFP (Supplementary Figure 7C), and the immunity-associated PM syntaxin, ROR2 (Figure 6), and instead, partially retained them in the ER. These observations indicate that CSEP0214 indeed hampers VPS18 activity.

An open question is how inhibition of this late step in the endomembrane trafficking pathway cause the marker proteins to accumulate in the ER. We propose this is due to congestion of the system, resulting in failure of efficient trafficking of cargo receptors, glycosyltransferases, proteases, chaperones and other proteins needed for the flow through the endomembrane trafficking pathway. Specific support for this idea is that CSEP0214 inhibits ST-mYFP from taking the first step from the ER to its destination in the Golgi apparatus despite the fact that VPS18 acts late in the pathway. A similar scenario appears to occur in protoplasts of the Arabidopsis *mon1* mutant, also arrested in a later step in this pathway. In that system, the two vacuolar proteins RD21-YFP and Aleu-GFP accumulate in enlarged MVBs and seemingly also in the ER (Cui et al., 2017).

The orthologous barley ROR2 and Arabidopsis PEN1 syntaxins are required for penetration resistance and for timely papilla formation at the site of powdery mildew fungal attack (Assaad et al., 2004; Böhlenius et al., 2010; Collins et al., 2003). They also recycle from the PM to accumulate in papillae, where they label EVs (Bhat et al., 2005; Böhlenius et al., 2010; Meyer et al., 2009; Nielsen et al., 2012; Nielsen et al., 2017). Interestingly, this recycling is dependent on the flippase ALA3 that also mediates recycling of PEN3, an ABC transporter likewise required for penetration resistance (Underwood et al., 2017). Notwithstanding this, we noted that penetration resistance is only marginally dependent on later steps in the endomembrane trafficking pathway (Liao et al., 2023; Nielsen et al., 2017). Thus, ESCRT-dependent MVB formation may not be directly involved in labelling papillae. Nevertheless, interference with VPS18 and VPS41 as well as expression of CSEP0214 hampered the accumulation of the ROR2 marker at the fungal attack sites. While it may be straightforward to envisage how *de novo* synthesized proteins become arrested in the ER due to congestion when a late step in the endomembrane trafficking pathway is interfered with, it is less obvious how ROR2 recycling is affected. Notably, cross-regulation exists between ESCRT-independent and ESCRT-dependent pathways in mammalian cells (Wei et al., 2021), and similar mechanisms may exist in plants affecting the ROR2 recycling, when the ESCRT-dependent endomembrane pathway is clogged.

Delivery of CSEP0214 affected encasement formation, *Mla1* and *Mla3*-mediated resistance, and HR (Figure 7). This is consistent with VPS18 interference and endomembrane pathway hampering, as encasement formation is dependent on the Rab5 GEF, VPS9a, as well as MON1 in Arabidopsis and barley (Nielsen et al., 2017; Liao et al., 2023). Also, CNL-mediated resistance requires a functional endomembrane pathway. Thus, HR activated by Arabidopsis RPM1 and RPS2 is dependent on the late endosome ESCRT components, AMSH3 and VPS4, in Arabidopsis and *N. benthamiana* (Schultz-Larsen et al., 2018), and barley powdery mildew resistance mediated by the CNL *Mla3* requires MON1, the target of CSEP0162 (Liao et al., 2023).

It is unclear how the endomembrane pathway contributes to CNL-induced HR, as CNLs oligomerize and form PM Ca^2+^ channels directly (Bi et al., 2021). However, two separate adaptor protein complexes, AP-2 and AP-4, involved in clathrin-coated vesicle formation, are required for RPM1 and RPS2-mediated resistance, while AP-4 in addition is required for tonoplast-PM fusion (Hatsugai et al., 2016; Hatsugai et al., 2018). We are looking forward to learn whether these adaptor protein functions link with the endomembrane pathway.

It is puzzling that CSEP0214 can suppress resistance by at least two *Mla* variants, and thus potentially target CNL-mediated resistance in general. Yet, even though *Bh* secretes these effectors, *Mla* resistance in barley is still functional. The existence of effectors with general resistance inhibitor functions, referred to as ‘silver bullet’ effectors, was previously questioned (Thordal-Christensen, 2020). However, this view may have to be revised. In fact, EDS1, which has a general and shared function in immunity mediated by Toll-interleukin-1 receptor (TIR) domain-containing NLRs (TNLs), has been described as an effector target (Bhattacharjee et al., 2011; Heidrich et al., 2011; Li et al., 2024; Wang et al., 2014). Thus, it appears that the barley powdery mildew fungus and other pathogens indeed secrete effectors that impede NLR functions in general. The puzzle is how NLRs can be fully effective despite the existence of such broadly acting effectors.

Additionally, we shed light on the mechanism of action of CSEP0214, and provide evidence that CSEP0214 blocks the function of VPS18 by interfering with its interaction with VPS16 (Figure 8). Both are core components of the HOPS and CORVET tethering complexes and VPS16, together with VPS33, form the SNARE-binding module (Baker and Hughson, 2016; Graham et al., 2013). Association of CSEP0214 with VPS18 may interfere with this SNARE-binding capacity, which would result in a failure of membrane fusion. The block could be achieved by direct attachment of CSEP0214 to the VPS16-binding surface in VPS18, which would prevent the interaction with VPS16. In this context it is interesting to note that the Y2H fragment of the VPS18 RING domain found in our Y2H cDNA screen covers most part of the binding area of VPS16 (Shvarev et al., 2022).

In conclusion, we have discovered that powdery mildew fungi share core effectors and one of them, CSEP0214, targets barley VPS18. We provide evidence that CSEP0214 inhibits the interaction of VPS18 and VPS16 of the endomembrane trafficking machinery transmitting material to the vacuole and for the secretion of EVs. Since this pathway is central for encasement formation and CNL-mediated immunity, we hypothesize that CSEP0214 makes a significant contribution to the virulence of *Bh*.

## Materials and methods

### In-silico analyses

The basis for the *B. hordei* (*Bh*) effector *in silico* analyses was the predicted secretome as defined by the 2018 annotation of the *Bh* genome (Frantzeskakis et al., 2018). Details of the bioinformatics analysis performed can be found in Supplementary File 1.

### Plant material and growth conditions

Barley (*Hordeum vulgare*) plants were grown in a climate chamber (16 h light (150 μE m^-2^ s^-1^)/8 h dark, at 20 °C, 60% relative humidity). Susceptible barley cultivar (cv.) Golden Promise seedlings were used for transient expression analysis, subcellular localization, callose staining, and RNAi experiments. Susceptible barley cv. Lottie was used in combination with the *Bh* isolate K1 for experiments to score host cell entry rates upon transient gene expression. Near-isogenic lines P01 and P02 of barley cv. Pallas (Kølster et al., 1986) were used to study resistance mediated by the *Mla1* and *Mla3* genes, respectively. *N. benthamiana* plants were used for Co-IP experiments and were grown in a controlled growth chamber (16 h light (150 μE m^-2^ s^-1^) at 23°C/8 h dark at 20°C, 60% relative humidity).

### Barley cDNA library

The used cDNA library was generated from leaf epidermal peels of two barley cultivars, inoculated with two different *Bh* isolates and sampled at six different time points prior to/after inoculation: Primary leaves of 6-to-7-day-old barley cv. Golden Promise seedlings were inoculated with *Bh* isolate DH14, while barley cv. Margret seedlings were inoculated with *Bh* isolate K1. Epidermal peels were collected at 0, 6, 12, 24, 48 and 96 hpi. Total RNA was extracted with the Qiagen RNeasy Plant Mini Kit (Qiagen, Hilden, Germany) and sent to ThermoFisher Scientific (Waltham, Massachusetts, USA) for cDNA library construction in pDEST22 (Y2H prey vector) based on oligo-dT-primed cDNA synthesis.

### Cloning

Barley and *Bh* cDNA coding sequences were amplified from RNA from infected barley plants, and cloned *via* Gateway^®^ BP reaction into pDONR201 or pDONR207 or by TOPO cloning reaction into pENTR/D-TOPO, according to the manufactureŕs protocol. Clones confirmed by sequencing were used for Gateway^®^ LR reaction into different destination vectors (see Supplementary Table 4).

### Gal4-based Y2H and split-ubiquitin Y2H experiments

Yeast cells were transformed following the standard LiAc-mediated protocol (Gietz and Woods, 2002). The heat shock at 42 °C was performed for 10 min for the PJ69-4A strain (James et al., 1996) (Gal4-based Y2H assays), and for 1 h for the THY.AP4 strain (Grefen et al., 2009) (split-ubiquitin Y2H experiments). Bait protein expression was verified via immunoblot with an α-Gal4 DBD antibody (Santa Cruz Biotechnology) or LexA antibody (gift by Karin Römisch), respectively. Yeast plasmid isolation was performed as described (Robzyk and Kassir, 1992). Yeast protein extraction was performed according to a protocol from the Dohlmann Lab (https://www.med.unc.edu/pharm/dohlmanlab/resources/lab-methods/tca/; (Cox et al., 1997)). The supernatant was used for the assessment of protein concentration by Bradford assay and immunoblot analysis.

### Co-Immunoprecipitation

*Agrobacterium tumefaciens* strain GV3101 (pMP90RK) was used for transient gene expression in leaves of *N. benthamiana*, which were harvested at 2 days after *A. tumefaciens* infiltration for protein extraction in 50 mM Tris-HCl pH 8.0, 1 mM EDTA, 1 mM dithiothreitol, supplemented with Complete Mini protease inhibitor cocktail (Roche, Mannheim, Germany)). Protein concentration was determined by the Bradford assay (Bradford, 1976), and 500 µg total protein was used for immunoprecipitation using α-RFP-Trap agarose beads (ChromoTek GmbH, Planegg, Germany). Samples of the soluble fraction (Input) and the bead-associated proteins (IP) were used for SDS-PAGE and immunoblot analysis using standard procedures. Details of antibodies used can be found in Supplementary Table 4.

### Immunoblot detection of CSEP0214

An antiserum was raised in rabbit against two CSEP0214-derived peptides (HTEGGKGGKGGKGDDDGDDD and IDAGCKSAADFACTCEHDTNKA), individually coupled to KLH carrier protein and designed to cover the predicted disordered region and the CFEM domain, respectively, and the resulting blood serum affinity-purified against the first peptide (Davids Biotechnologie GmbH, (Regenburg, Germany). Total protein extract from non-infected and *Bh*-infected barley leaves sampled at 5 days post inoculation (dpi) were used for immunoblot analysis. For detection of purified CSEP0214 protein from *E. coli*, the Rosetta strain was transformed with pDEST15-RFP or pDEST17-CSEP0214 protein expression constructs, induced with 1 mM isopropyl-β-D-thiogalactopyranoside, and incubated at 16 °C overnight. Cell extract was obtained by sonication, which was further used for SDS-PAGE and immunoblot analysis.

### Transient expression of gene constructs in barley

Transformation of gene constructs into leaf epidermal cells of the abaxial side of 7-day-old barley seedlings was conducted by particle bombardment as described before (Douchkov et al., 2005) using 1 µm gold particles *via* the Bio-Rad PDS-1000/He particle delivery system, mounted with a hepta-adapter, according to the manufacturer’s instructions (Bio-Rad). Fluorescent signals emitted by mCherry, RFP or mYFP were visualized using a Leica (Wetzlar, Germany) Stellaris 5 confocal laser scanning microscope. See Supplementary Table 4 for a list of the marker constructs used in this work.

### Study of immune responses

Papilla formation, penetration resistance and encasement formation were studied after transient transformation of barley leaf epidermal cells with the respective constructs in combination with a construct for β-glucuronidase (GUS) expression (Douchkov et al., 2005), followed by inoculation with *Bh*, and Aniline blue staining, which was subsequently assessed by UV epifluorescence microscopy. Papilla formation was studied at 1 dpi whereas penetration resistance was studied at 2 dpi. Encasement formation was studied at 4 dpi after bombardment and tetraconazole treatment (100 μg/ml tetraconazole in 20% acetone with 0.04% Tween-20 (Maffi et al., 1995)). To study HR and fungal development, the EtHAn strain of *P. fluorescence* (Thomas et al., 2009) was used, followed by inoculation with avirulent *Bh* isolates C15 and A6, respectively. Cells undergoing HR were stained with trypan blue as described before (Koch and Slusarenko, 1990) and fungal structures were visualized by Coomassie staining as above. Scoring was performed by light microscopy at 4 dpi.

## Abbreviations

3-AT: 3-amino-1,2,4-triazole
ARF: ADP ribosylation factor
Bh: Blumeria hordei
Bgt: *Blumeria graminis* f.sp. *tritici*
CFEM: common in fungal extracellular membrane
CNL: coiled-coil, nucleotide-binding, leucine-rich repeat
Co-IP: co-immunoprecipitation
CORVET: class C core vacuole/endosome tethering
CSEP: candidate for secreted effector protein
cv.: cultivar
dpi: days post inoculation
EHM: extrahaustorial membrane
ER: endoplasmic reticulum
ESCRT: endosomal sorting complex required for transport
EV: extracellular vesicles
FL: full-length
GUS: β-glucuronidase
HA: hemagglutinin
HOPS: homotypic fusion and protein-sorting
hpi: hours post infection
HR: hypersensitive response
MVB: multivesicular body
PAMP: pathogen-associated molecular pattern
PM: plasma membrane
PTI: pattern-triggered immunity
RING: really interesting new gene
RNAi: RNA interference
SDS: sodium dodecyl sulfate
SNARE: *N*-ethylmaleimide-sensitive-factor attachment receptor
SP: signal peptide
TPM: transcripts per million
VPS: vacuolar protein sorting
Y2H: yeast two-hybrid

## Author contributions

BS performed the *in silico* CSEP analysis and conducted all protein-protein interaction assays. SD performed all cell biological (co-)localization and immune response studies.

HTC and RP conceived the study and supervised the project.

PDS and RP designed the Y2H prey library and coordinated their synthesis. BS (with the help of others) collected material for the Y2H library.

BS and SD drafted the manuscript; PDS, HTC and RP edited the manuscript.

## Acknowledgements

We acknowledge the support of many helping hands in the Panstruga and Spanu labs (amongst others Lamprinos Frantzeskakis, Hannah Kuhn, Anja Reinstädler, Hongpo Wu, Linhan Li, and Helen Pennington) regarding the isolation of epidermal peels for cDNA construction. We thank our late and good friend, Dr. Patrick Schweizer (Leibniz Institute of Plant Genetics and Crop Plant Research, Gatersleben, Germany) for providing the RNAi vector. We appreciate the cloning of the pAMpat-SP-mCherry construct by Mark Kwaaitaal and Meike Lauts. We thank the Centre for Advanced Bioimaging (CAB) at the University of Copenhagen for use of their facilities. Furthermore, we thank Prof. Chris Hawes for making the ST–YFP construct available to us. We thank Prof. Andreas Nebenführ for the SP–mCherry–HDEL construct. We thank Prof. Jeff Chang for providing the EtHAn bacterial strain and Prof. Jonathan Jones for allowing us to use the pEDV6 vector. We thank the late Dr. Patrick Schweizer (Leibniz Institute of Plant Genetics and Crop Plant Research, Gatersleben, Germany) for providing the RNAi vector and Karin Römisch (Saarland University, Germany) for sharing the LexA antibody with us.

## Supplementary material

**Supplementary Figure 1.**
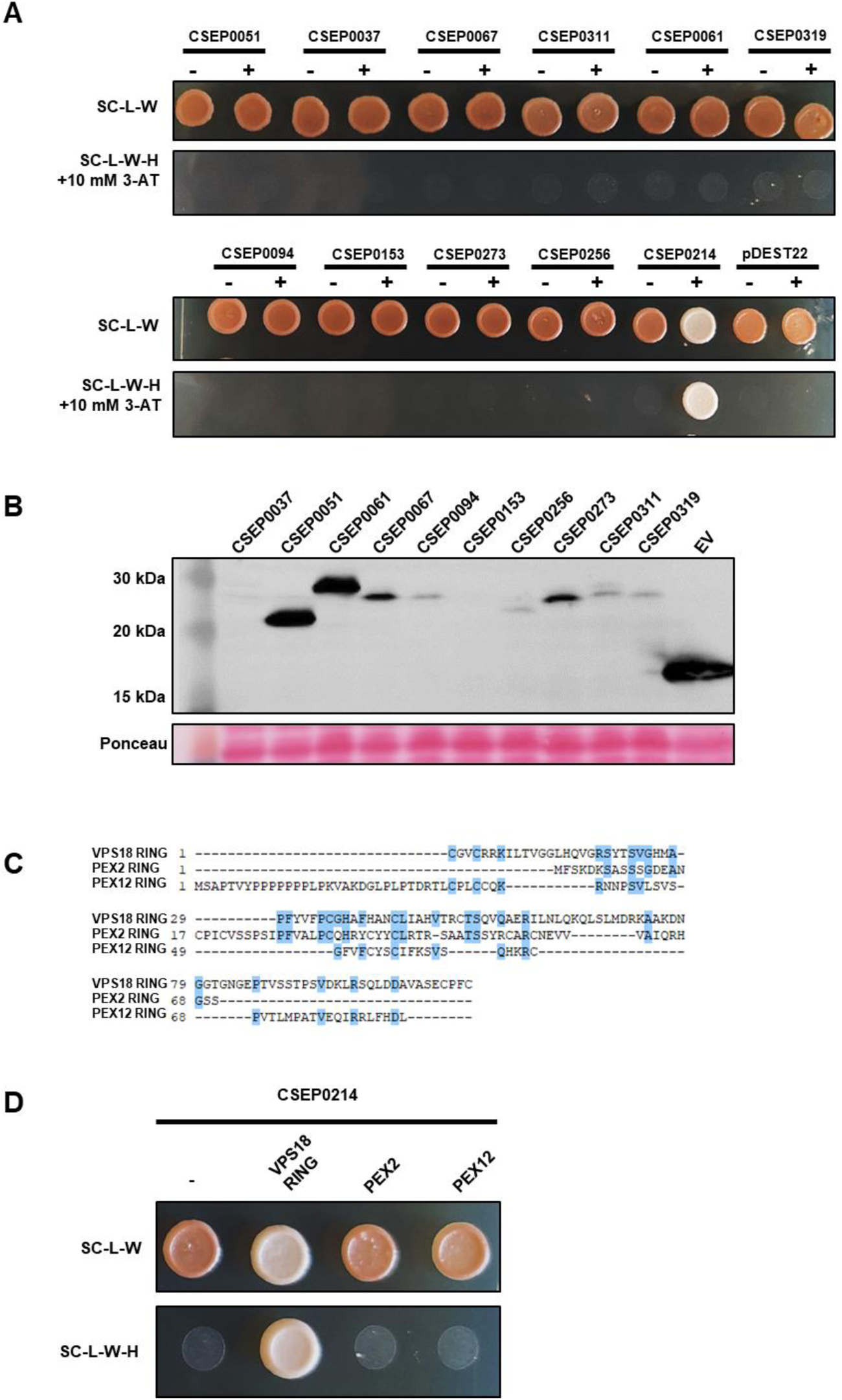
Specificity of the interaction between CSEP0214 and VPS18. **A.** Gal4-based Y2H assay of the VPS18 RING domain fused to the Gal4 activation domain (+) was tested against ten randomly selected CSEPs fused to the Gal4 binding domain. The empty Gal4 activation domain vector (pDEST22) (-) was used as negative control, CSEP0214 as positive control. Shown on the top of each panel is yeast growth on medium that is selective for the presence of the plasmids (SC-L-W; growth control). Shown on the bottom of each panel is yeast growth on interaction-selective medium (SC-L-W-H+10 mM 3AT). **B**. The expression of the different CSEPs that do not show interaction with VPS18 RING was verified in yeast cells by immunoblot analysis with an antibody against the Gal4 binding domain. On the left, molecular masses of marker proteins are given. Ponceau staining was used to demonstrate equal amounts of total proteins on the blots. Expected molecular weights of the respective proteins: BD-CSEP0037 (35 kDa), BD-CSEP0051 (26 kDa), BD-CSEP0061 (32 kDa), BD-CSEP0067 (34 kDa), BD-CSEP0094 (30 kDa), BD-CSEP0153 (34 kDa), BD-CSEP0256 (29 kDa), BD-CSEP273 (32 kDa), BD-CSEP0311 (30 kDa), BD-CSEP0319 (32 kDa). **C**. Amino acid sequence alignment of the RING domains of barley VPS18 with the RING domains of two unrelated barley peroxisomal proteins, PEX2 and PEX12. Conserved (>60%) amino acids are highlighted in blue. **D.** Gal4-based Y2H assay of the PEX2 and PEX12 RING domains (fused to the Gal4 activation domain) with CSEP0214 (fused to the Gal4 binding domain). The empty pDEST32 vector (-) was used as negative control and the VPS18 RING domain (+) was used as positive control. Shown on the top is yeast growth on medium that is selective for the presence of the plasmids (SC-L-W; growth control). Shown on the bottom is yeast growth on interaction-selective medium (SC-L-W-H). Experiments were performed three times with similar results. Images were taken after 3 days of growth.

**Supplementary Figure 2.**
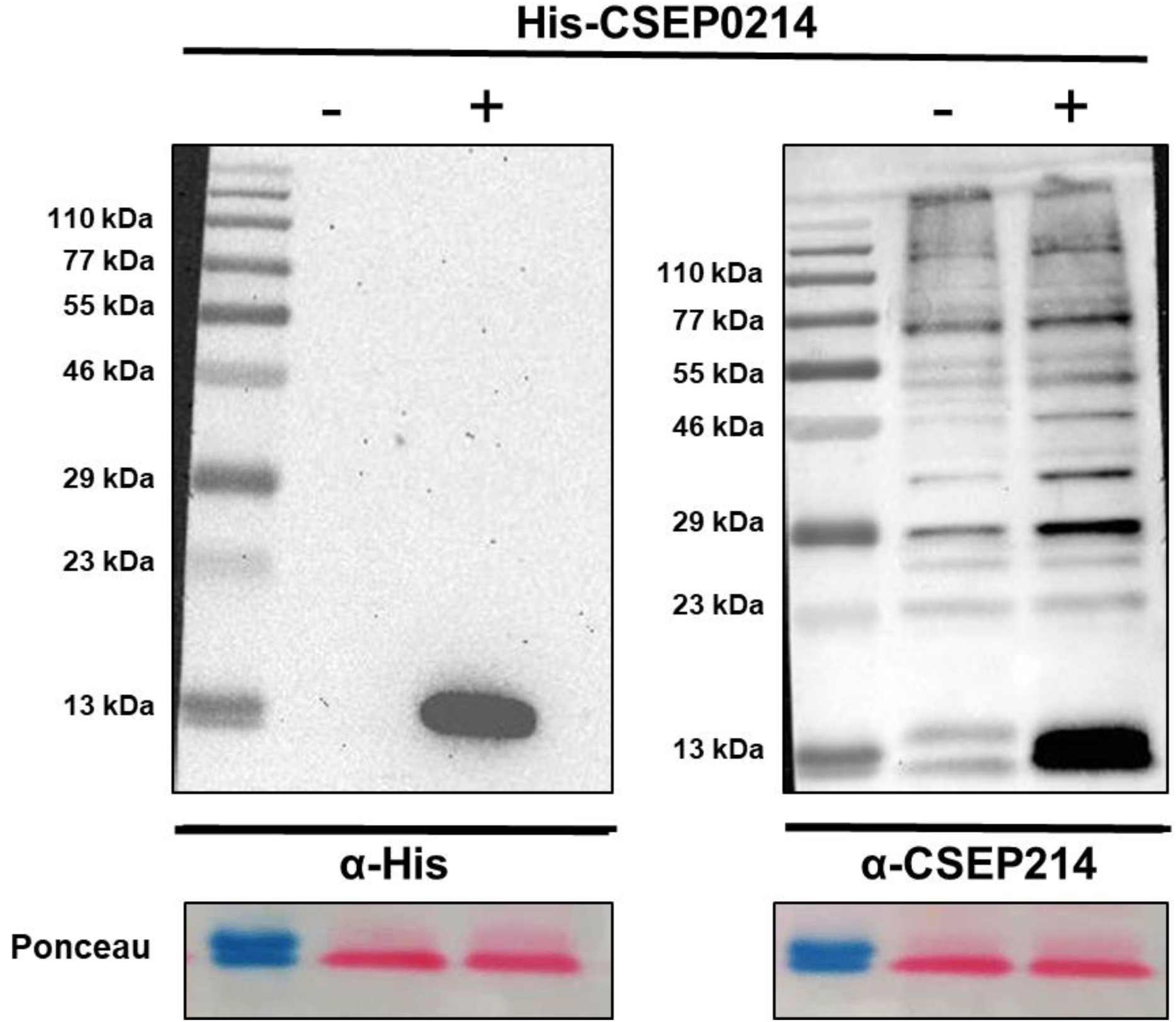
Test of the α-CSEP0214 antibody with heterologously expressed His-CSEP0214 from *E. coli*. *E. coli* Rosetta strains carrying pDEST15-RFP (-) as negative control, and pDEST17-CSEP0214 (+) were used to generate protein extracts for SDS-PAGE and immunoblot analysis. The membranes were probed with α-His and α-CSEP0214 antibodies, respectively. On the left, molecular masses of marker proteins are given. Ponceau staining was used to demonstrate equal amount of total proteins on the blots.

**Supplementary Figure 3.**
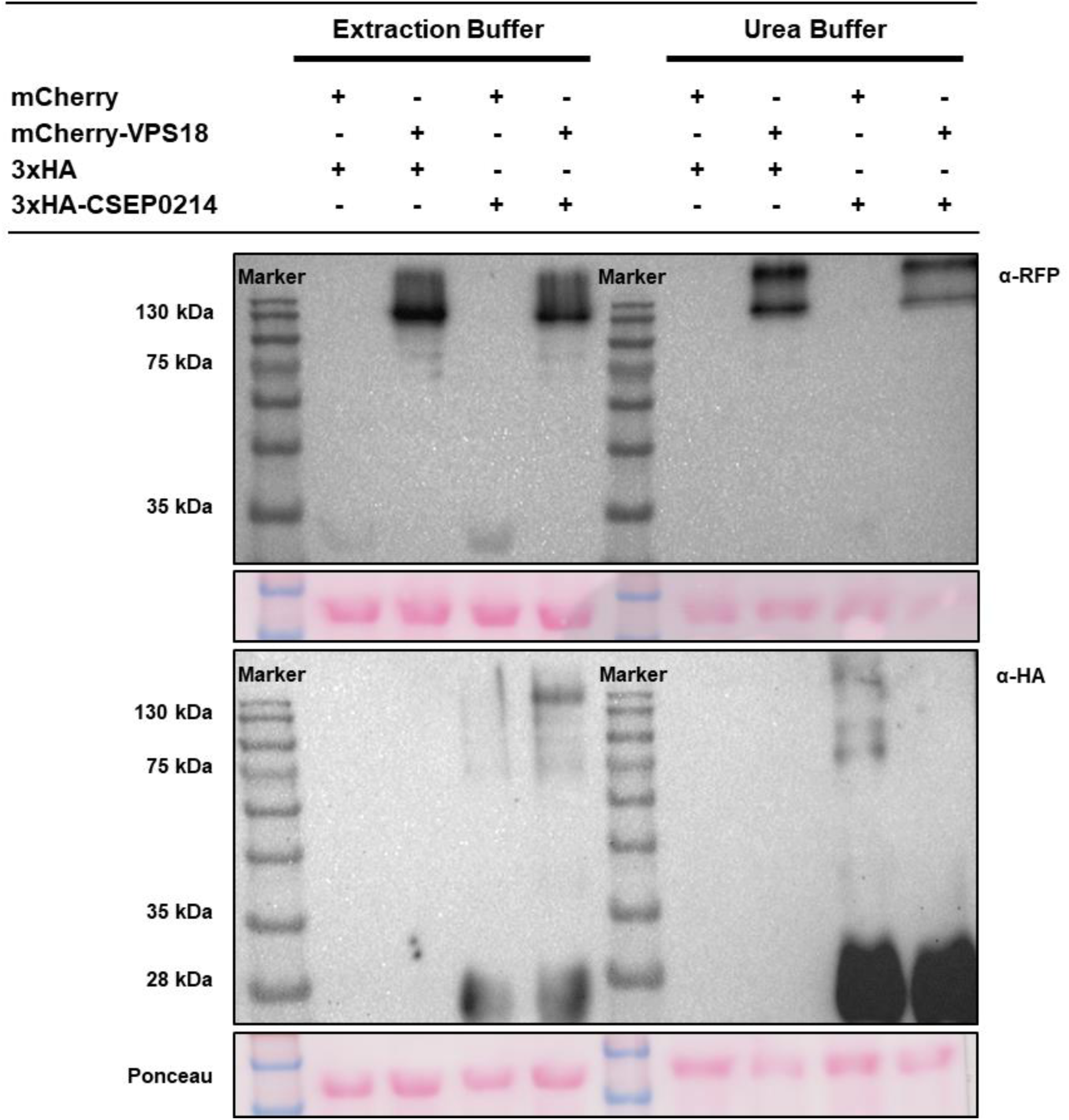
CSEP0214 and VPS18 form a stable, heat-resistant protein complex. Fusion proteins mCherry-VPS18 and 3xHA-CSEP0214 or the mCherry and 3xHA tags alone were co-expressed in different combinations via *A. tumefaciens*-mediated transient gene expression in leaves of *N. benthamiana* plants. Total protein extraction was performed at 3 days after agroinfiltration with either a standard extraction buffer (10 mM Tris-HCl pH 8, 1 mM EDTA, 1 mM dithiotreitol, protease inhibitor cocktail) or a denaturing-urea buffer (7.7 M Urea, 2 M thiourea, 300 mM NaCl, 0.25% CHAPS, 50 mM NaH_2_PO_4_ pH 8, 50 mM Tris pH 8). Protein extracts were used for SDS-PAGE and immunoblot analysis. The membranes were probed with *α-*RFP and *α-*HA antibodies, respectively. On the left, molecular masses of marker proteins are given. Ponceau staining was used to demonstrate equal amount of total proteins on the blots.

**Supplementary Figure 4.**
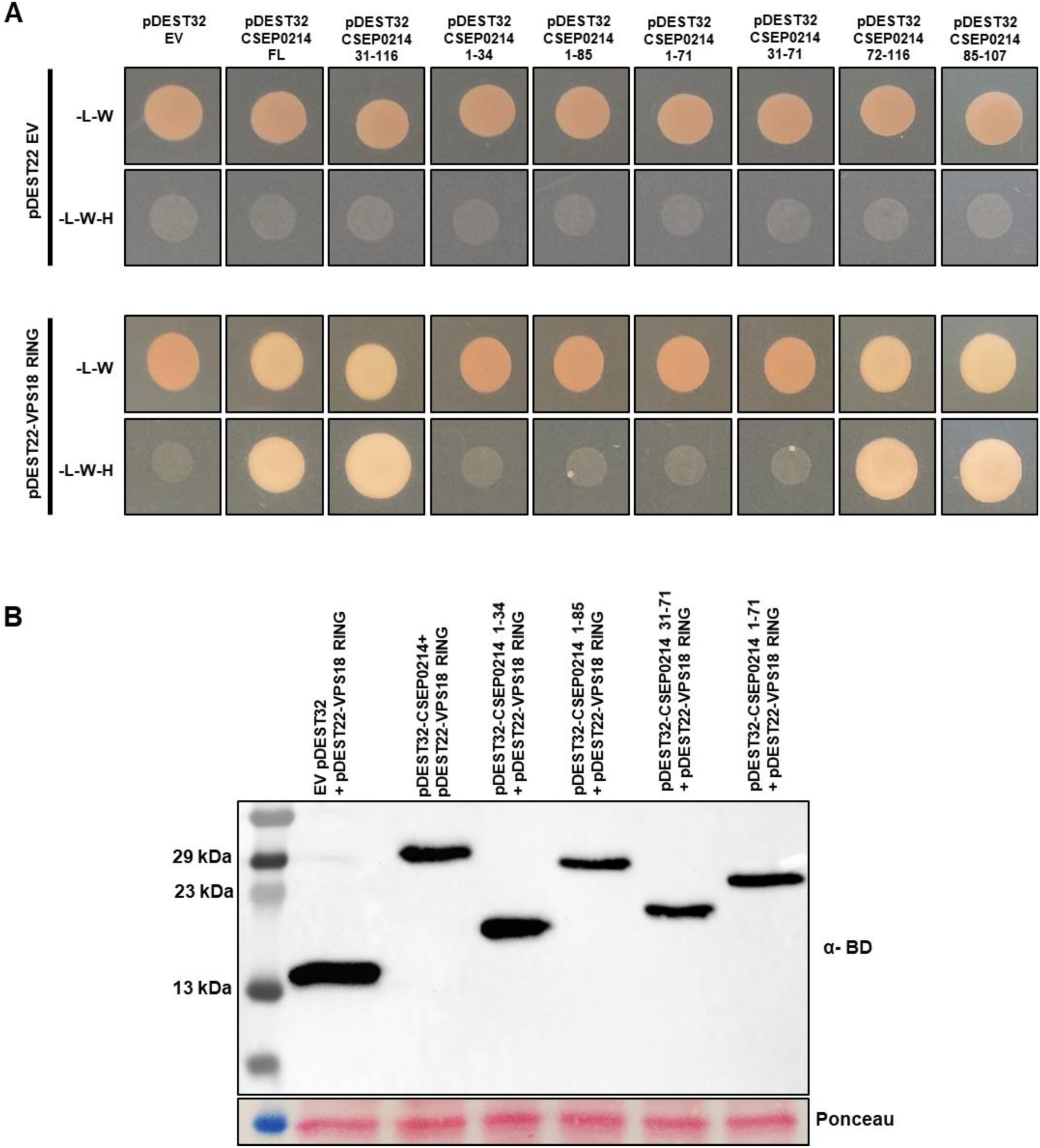
Y2H assay of the VPS18 RING with truncations of CSEP0214. **A.** Gal4-based Y2H assay of the VPS18 RING domain fused to the Gal4 activation domain was tested against truncations of CSEP0214 (see Fig. 4A) fused to the Gal4 binding domain. The empty Gal4 activation domain vector (pDEST22) was used as negative control, CSEP0214 full length (FL) as positive control. Shown on the top of each panel is yeast growth on medium that is selective for the presence of the plasmids (-L-W; growth control). Shown on the bottom of each panel is yeast growth on interaction-selective medium (-L-W-H). **B**. The expression of the CSEP0214 truncations that do not show interaction with VPS18 RING was verified in yeast cells by immunoblot analysis with an antibody against the Gal4 binding domain. On the left, molecular masses of marker proteins are given. Ponceau staining was used to demonstrate equal amounts of total proteins on the blots. Expected molecular weights of the respective proteins: pDEST32 EV (16 kDa), BD-CSEP0214 (30 kDa), BD-CSEP0214 1-34 (19 kDa), BD-CSEP0214 1-85 (26 kDa), BD-CSEP0214 31-71 (20 kDa), BD-CSEP0214 1-71 (23 kDa).

**Supplementary Figure 5.**
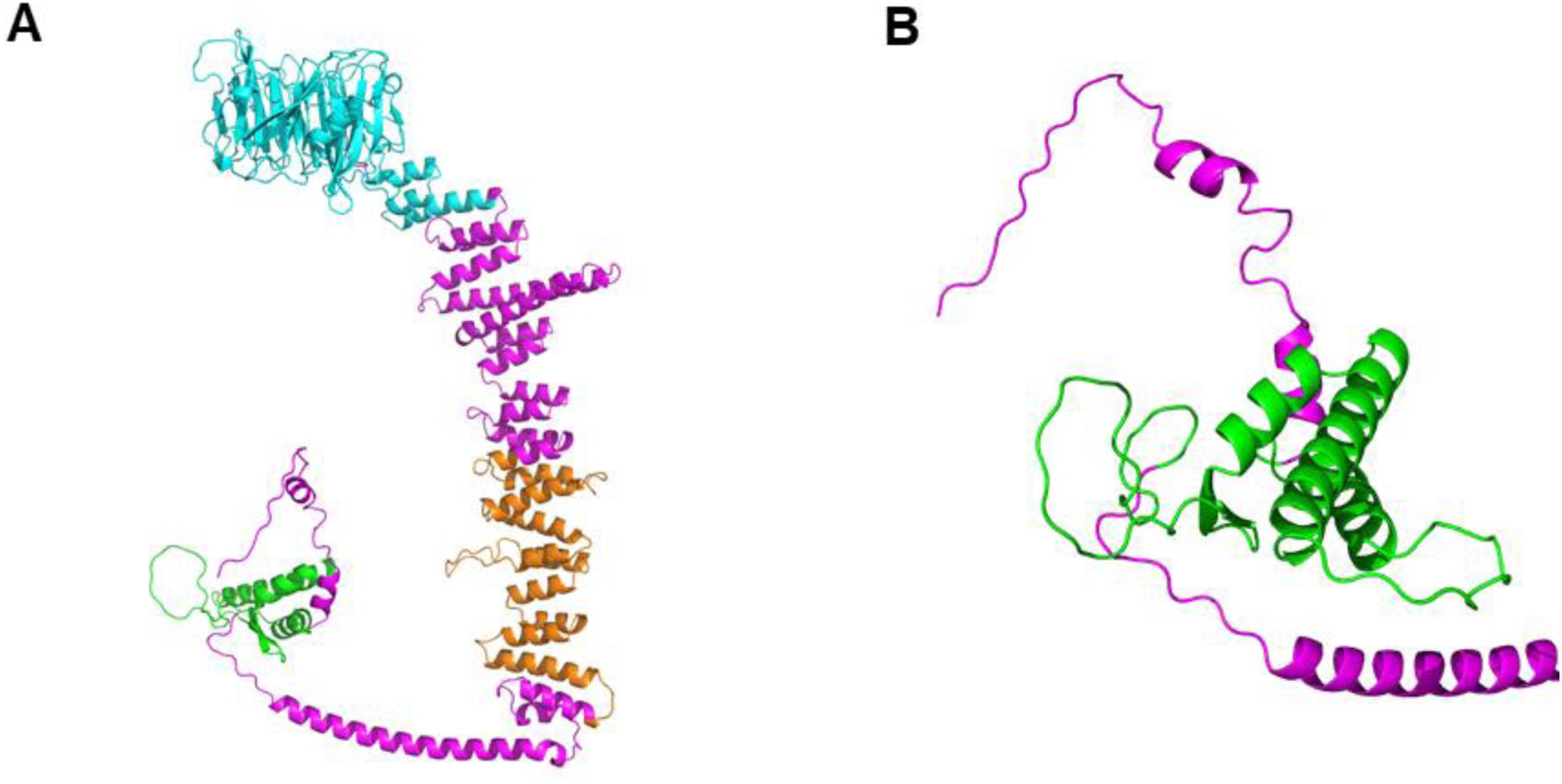
Predicted three-dimensional structure of barley VPS18. **A**. Predicted Alphafold2 structure of barley VPS18. Predicted domains are indicated as follows: Pep3/VPS18/beta-propeller domain (blue), Clathrin, heavy chain/VPS, 7-fold repeat (orange) and Zinc finger, RING-type domain (green), and undefined regions (magenta). **B**. Magnification of the RING domain, which includes the potential interaction site with CSEP0214.

**Supplementary Figure 6.**
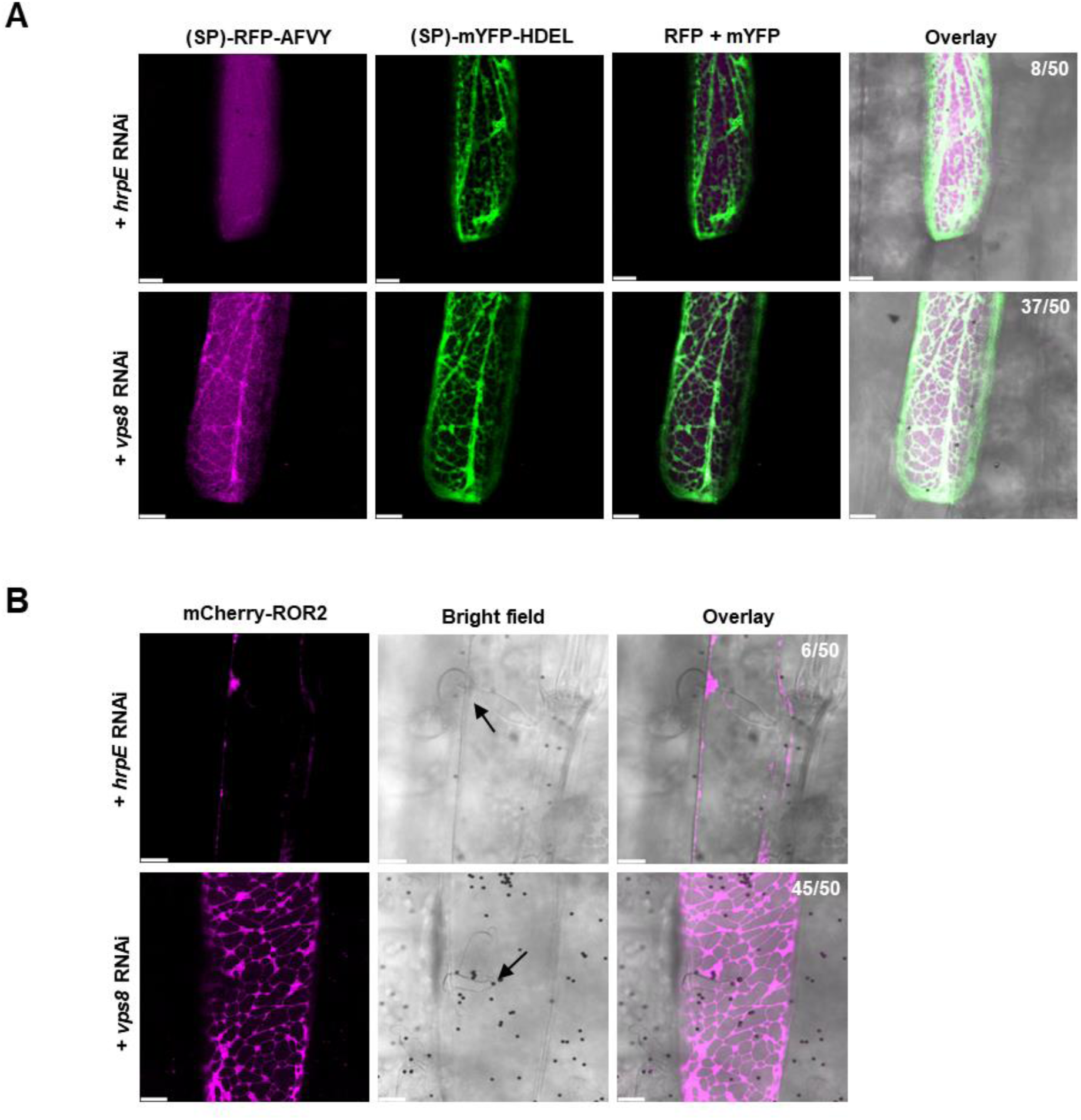
Transient knockdown of *VPS8* in barley leaf epidermal cells perturbs the subcellular localization of vacuolar and papilla markers. The vacuolar marker, (SP)-RFP-AFVY and the ER marker (SP)-mYFP-HDEL (**A**), or mCherry-ROR2 (**B**), were co-expressed with either the *hrpE* control RNAi construct (pIPKTA30N::*hrpE*) or knockdown construct of *VPS8* (pIPKTA30N::*VPS8*) in barley (cv. Golden Promise) leaf epidermal cells. mCherry-ROR2 was observed at the papilla 24 hpi with *Bh* isolate C15. Numbers in the top right corner of the overlay panels indicate the occurrence of unusual marker localization in 50 inspected cells. Scale bar, 10 µm.

**Supplementary Figure 7.**
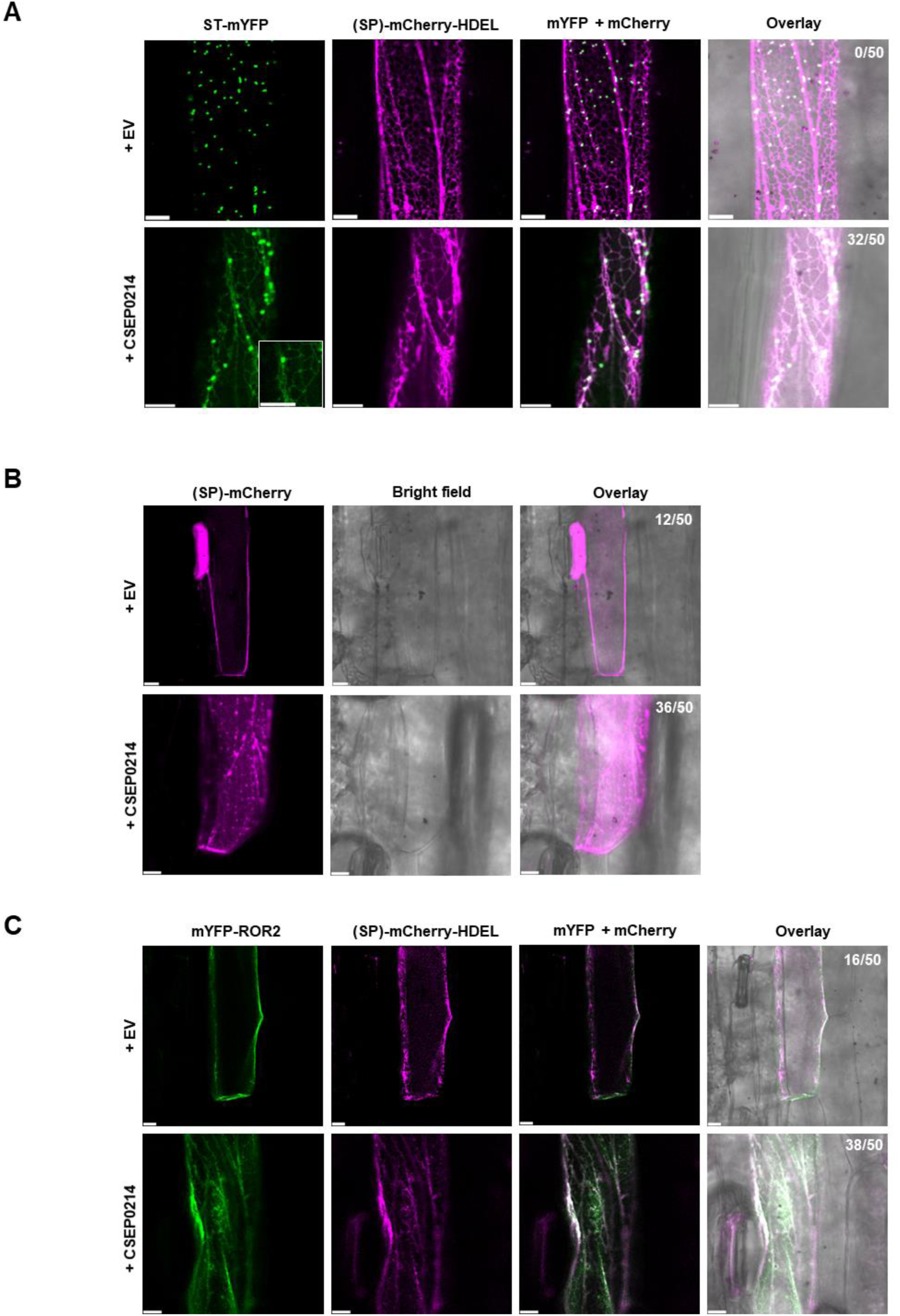
Expression of CSEP0214 in barley leaf epidermal cells perturbs the subcellular localization of markers of the endomembrane trafficking pathway. The Golgi marker ST-mYFP and the ER marker (SP)-mCherry-HDEL (**A**), secreted marker protein (SP)-mCherry (**B**), or the mCherry-ROR2 and the ER marker (SP)-mYFP-HDEL (**C**), were co-expressed with either an empty vector (EV; pUbi::GW) or expression of CSEP0214 (pUbi::CSEP0214) in barley (cv. Golden Promise) leaf epidermal cells. Inset (**A**), ST-mYFP appears to be in the ER network. Numbers in the top right corner of the overlay panels indicate the occurrence of unusual marker localization in 50 inspected cells. Scale bar, 10 µm.

**Supplementary Figure 8.**
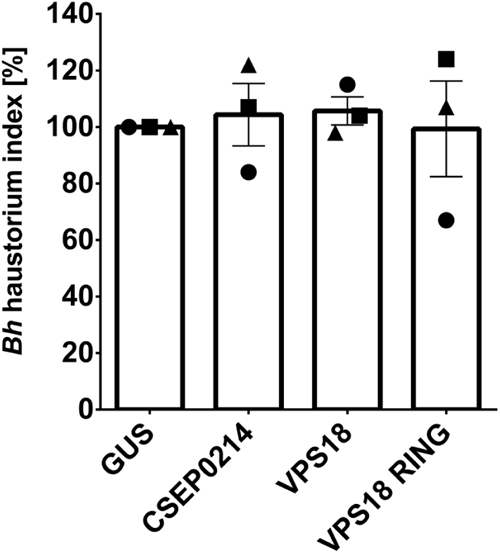
Expression of CSEP0214, VPS18 or VPS18 RING does not affect the fungal penetration rate. Leaf epidermal cells of seven-day-old barley plants (cv. Lottie) were transformed by particle bombardment with a construct expressing GUS along with either an empty vector (EV; pUbi::GW) or expression of CSEP0214 (pUbi::CSEP0214), VPS18 (pUbi::VPS18) or VPS18 RING (pUbi::VPS18 RING). After one day, the leaves were inoculated with *Bh* isolate K1 and at 48 hpi the percentage of GUS cells with haustoria was quantified. n = 3. Error bars, SE. Data assessed by Student’s T-tests.

