## Supplementary File 1 for "A powdery mildew core effector protein targets host endosome tethering complexes HOPS and CORVET"

***In-silico* analyses of *B. hordei* secreted proteins**

The Galaxy server (Galaxy Community, 2022) was used to perform a TBLASTN search against the mentioned powdery mildew fungal genomes (Kusch et al., 2023) by constructing a custom BLAST library of 17 powdery mildew genomes (*B. graminis* f.sp. *tritici* 96224, *B. graminis* f.sp. *triticale* THUN12, *Erysiphe alphitoides* MS-42D*, Erysiphe necator* C, *Erysiphe neolycopersici* UMSG2, *Erysiphe pisi* Palampur-1, *Golovinomyces cichoracearum* UCSC1*, Golovinomyces cichoracearum* UMSG3, *Golovinomyces magnicellulatus* FPH2017-1*, Golovinomyces orontii* MGH*, Leveillula taurica* HNHM-MZC-006405, *Parauncinula polyspora*, *Podosphaera xanthii* 2086*, Podosphaera xanthii* Wanju, *Phyllactinia moricola* HMJAU-PM91933, *Pleochaeta shiraiana* HAL3440 F, and *Podosphaera leucotricha* PuE-3)*.* Amino acid sequences, encoded by 15 out of the 16 genomes were filtered subsequently via TBLASTN searches against the genomes of non-phytopathogenic (*Aspergillus* spp., *Penicillium* spp., and *Saccharomyces* spp.) and phytopathogenic (*Alternaria* spp., *Bipolaris* spp., *Botrytis* spp., *Claviceps* spp., *Colletotrichum* spp., *Curvularia* spp., *Drechslera* spp., *Drepanopeziza* spp., *Exserohilum* spp., *Fusarium* spp., *Gaeumannomyces* spp., *Magnaporthe* spp., *Monilinia* spp., *Pyrenophora* spp., *Pyricularia* spp., *Ramularia* spp., *Rhynchosporium* spp., *Sclerotinia* spp., *Septoria* spp., *Thielaviopsis* spp., *Venturia* spp., *Verticillium* spp., *Zymoseptoria* spp.) ascomycotes in the NCBI database with a cut-off e-value of e^-10^. Unique powdery mildew secreted proteins are considered as core effectors. SignalP 4.1 (Petersen et al., 2011), SignalP 5.0 (Almagro Armenteros et al., 2019), SignalP 6.0 (Teufel et al., 2022) and Phobius (Käll et al., 2004) were used on the set of 800 secreted proteins for prediction of SP and TMHMM 2.0 (Krogh et al., 2001) was subsequently used for prediction of TM domains. The phylogenetic tree of all 800 secreted proteins of *Bh* was established with Mega11 (Tamura et al., 2021) by aligning all amino acid sequences with the MUSCLE algorithm and using the Maximum Likelihood method and JTT matrix-based model (Jones et al., 1992). The consensus tree was inferred from 1,000 bootstrap replicates (Felsenstein, 1985). For the phylogenetic tree of the powdery mildew fungi, all depicted genomes were analyzed with BUSCO (Waterhouse et al., 2018) to identify single-copy genes. These genes were used in Orthofinder (Emms and Kelly, 2019) for further analysis and to construct the tree with the STAG algorithm. The prediction of potential effector proteins and their localization was accomplished with EffectorP 1.0 (Sperschneider et al., 2016), EffectorP 2.0 (Sperschneider et al., 2018a), EffectorP 3.0 (Sperschneider and Dodds, 2022), LOCALIZER (Sperschneider et al., 2017) and ApoplastP (Sperschneider et al., 2018b). The InterPro (Jones et al., 2014) library was downloaded and run locally to identify known protein domains. To complement this approach, the Phyre^2^ server (Kelley et al., 2015) was utilized to model each protein structure and to predict functions based on known proteins. Further, the 3D structures of all 800 proteins were modelled with Alphafold2 (Jumper et al., 2021) to perform structural searches. Published RNA-seq datasets were downloaded from the NCBI Sequence Read Archive (<https://www.ncbi.nlm.nih.gov/sra>), accessions PRJNA184807 (Hacquard et al., 2013), PRJNA325323 (Lu et al., 2016), PRJNA393407 (Sucher et al., 2018), PRJNA416115 (Frantzeskakis et al., 2018), PRJNA433341 (Saur et al., 2019), PRJNA835302 (Qian et al., 2023) and PRJNA639160 (Kusch et al., 2021). The data were adapter- and quality-trimmed using Trimmomatic v0.39 (Bolger et al., 2014) using the options 'SLIDINGWINDOW:3:18 LEADING:6 TRAILING:6 MINLEN:70’. Read quality was confirmed via FastQC v0.11.5 (Babraham Bioinformatics, Cambridge, UK). Reads were mapped to the *Bh* DH14 v4 reference genome (Frantzeskakis et al., 2018) with HISAT2 v2.0.5 (Kim et al., 2015), read count tables generated via featureCounts v2.0.1 (Liao et al., 2014), and TPM values calculated in R v4.1.2 (R Core Team, 2021). To compare the gene models of the annotations of 2010 and 2018, respectively, the CDS of the CSEPs were searched by BLAST against the new gene models and against the *Bh* genome.
